# New insights into the earlier evolutionary history of epiphytic macrolichens

**DOI:** 10.1101/2021.08.02.454570

**Authors:** Qiuxia Yang, Yanyan Wang, Robert Lücking, H. Thorsten Lumbsch, Xin Wang, Zhenyong Du, Yunkang Chen, Ming Bai, Dong Ren, Jiangchun Wei, Hu Li, Yongjie Wang, Xinli Wei

## Abstract

Lichens are well known as pioneer organisms colonizing bare surfaces such as rocks and therefore have been hypothesized to play a role in the early formation of terrestrial ecosystems. Given the rarity of fossil evidence, our understanding of the evolutionary history of lichen-forming fungi is primarily based on molecular dating approaches. These studies suggest extant clades of macrolichens diversified after the K–Pg boundary. Here we corroborate the mid-Mesozoic fossil Daohugouthallus ciliiferus as an epiphytic macrolichen that predates the K-Pg boundary by 100 Mys. Based on new material and geometric morphometric analysis, we demonstrate that the Jurassic fossil is morphologically most similar to Parmeliaceae, but cannot be placed in Parmeliaceae or other similar family-level clades forming macrolichens as these evolved much later. Consequently, a new family, Daohugouthallaceae, is proposed here to accommodate this fossil, which reveals macrolichens may have been diverse long before the Cenozoic diversification of extant lineages.

## Introduction

Lichens are a stable symbiosis composed of fungi and algae and/or cyanobacteria; they also include a diverse microbiome (*Spribille et al., 2016*; *Lücking and Nelsen, 2018*; *Hawksworth and Grube, 2020*). Lichens are components of mostly terrestrial ecosystems from the polar regions to the tropics *(Lumbsch and Rikkinen, 2017*), growing on all kinds of substrates, including bark, rock, leaves and soil (*Belnap et al., 2001*; *Brodo et al., 2001*; *Nash, 2008*). Lichens play important roles in ecosystem function, including weathering of rock and accelerating formation of soil (*Lindsay, 1978*; *Chen et al., 2000*), fixing carbon and nitrogen from the atmosphere (*Wu et al., 2011*), and as food source for animals (including humans, *Cornelissen et al., 2007*). Due to their sensitivity to environmental changes, lichens also are widely used as bioindicator of air pollution and environmental health. The evolutionary history of lichen-forming fungi is poorly understood, due to the sparse fossil record, and has been primarily reconstructed based on molecular dating analyses (*Kraichak et al., 2015*, *2018*; *Lücking et al., 2017*; *Huang et al., 2019*; *Widhelm et al., 2019*; *Nelsen et al., 2020*). Although these approaches proposed a framework to illustrate how the lichen symbiosis may have evolved, the fossil evidence is indispensable in testing and supplementing the current understandings especially when the earlier fossil was discovered.

To date, about 190 fossils are accepted to represent genuine lichens, and a few are considered ambiguous, i.e. potentially representing lichens (*Lücking and Nelsen, 2018*). The earliest accepted lichen are two crustose lichens from Devonian fossils, i.e. Cyanolichenomycites devonicus and Chlorolichenomycites salopensis (419–411 Mya), which were inferred to be saxicolous or terricolous (*Lücking and Nelsen, 2018*). The other earlier lichen is from the Lower Cretaceous, i.e. *Honeggeriella complexa* (ca. 133 Mya) that was suggested as a squamulose or foliose lichen, although no larger pieces showing its architecture are preserved (*Matsunaga et al., 2013*; *Honegger et al., 2013*). A recent study corroborated the lichen property of *Daohugouthallus ciliiferus* from the Middle Jurassic (ca. 165 Mya) that represented the earliest foliose to subfruticose lichen, based on its external morphology (*Fang et al., 2020*). Other than this fossil, lichens with foliose and fruticose thalli, the principle forms of macrolichens, have no unambiguous fossil record before the K–Pg boundary (*Lücking and Nelsen, 2018*), and the oldest accepted macrolichen fossils are from Eocene Baltic amber (38–44 Mya), including foliose and fruticose Parmeliaceae (*Kaasalainen et al., 2015*, *2017*). *Ziegler (2001)* described and depicted various fragments of presumed parmelioid macrolichens from the Keuper formation (230–200 Mya), a work that has not received much attention and was overlooked by lichenologists (*Lücking and Nelsen, 2018*). A revision of the data presented in that study did not show conclusive evidence for the presence of macrolichens in the depicted fossils; it also presents methodological issues, such as lack of voucher information and precise links of micrographs to the corresponding fossils. Given this lack of evidence for macrolichen fossils prior to the K–Pg boundary, the question arises about the significance of the Jurassic lichen *Daohugouthallus ciliiferus* for understanding the evolutionary history of macrolichens in the Mesozoic.

Macrolichens evolved independently in various classes of Basidiomycota (Agaricomycetes: e.g., *Cora*) and Ascomycota, such as Arthoniomycetes (e.g., *Roccella*) and Lichinomycetes (e.g., *Thyrea*).

However, most macrolichens are found in the largest class of lichenized Ascomycota, the Lecanoromycetes. According to recent molecular clock studies (*Nelsen et al., 2020*), within Lecanoromycetes, macrolichens evolved independently in at least three separate clades: 1) Umbilicariaceae (subclass Umbilicariomycetidae), with an inferred stem age of 164–150 Mya; 2) Icmadophilaceae (subclass Ostropomycetidae), including macrolichens such as *Siphula* and *Thamnolia*, with an inferred stem age of 214–190 Mya; and 3) subclass Lecanoromycetidae, which contain major macrolichen clades such as Cladoniaceae, Coccocarpiaceae, Collemataceae, Pannariaceae, Parmeliaceae, Peltigeraceae, Physciaceae, Ramalinaceae, and Teloschistaceae. Among these, Collemataceae appears to be the oldest, with a stem age of around 145–140 Mya, followed by Pannariaceae and Coccocarpiaceae (140-135 Mya), Peltigeraceae (125-120 Mya), Parmeliaceae (80–75 Mya), Physciaceae (80–60 Mya for macrolichen clades), and Cladoniaceae (60–55 Mya), Teloschistaceae (65–35 Mya for macrolichen clades) and Ramalinaceae (around 50 Mya for macrolichen clades; *Nelsen et al. 2020*). However, even families that appeared in the Cretaceous, such as Parmeliaceae and Peltigeraceae, were shown to exhibit significantly increased diversification rates after the Cretaceous–Paleogene (K–Pg) boundary (66 Mya, *Kraichak et al., 2018*; *Huang et al., 2019*; *Widhelm et al., 2019*; *Nelsen et al., 2020*).

Given that macrolichens have evolved in convergent fashion in multiple, often unrelated lineages in Asco- and Basidiomycota, it is vital to clarify the position of *Daohugouthallus ciliiferus* in reconstructing and understanding the evolution of macrolichens. The oldest known macrolichen fossil, *Daohugouthallus ciliiferus*, cannot be placed with certainty in extant lineages because fine-scaled diagnostic features, such as hamathecium, ascus, and ascospore structure, cannot be observed. However, techniques such as automated image recognition have allowed to at least analyze morphometric features in a way that allow a quantitative approach to hypothesis testing. In the present study, we therefore provide an updated morphological assessment of *Daohugouthallus ciliiferus*, using an image-based, geometric morphometric analysis to compare the fossil with a range of extant macrolichens. In parallel, we used the large molecular clock analysis by *Nelsen et al. (2020)*, which due to the comprehensive sampling offers a much broader framework than other molecular clock studies including lichen formers (e.g. *James et al., 2006*; *Lutzoni et al., 2018*; *Kraichak et al., 2018*), to reassess stem and crown node ages for major clades of macrolichen formers in the Lecanoromycetes, in comparison to the age of the fossil. As a result, we propose a new family, Daohugouthallaceae, for this fossil.

## Results

### Daohugouthallaceae

X.L. Wei, X. Wang, D. Ren & J.C. Wei, fam. nov. (*Fig. 1*)

**Fig. 1.**
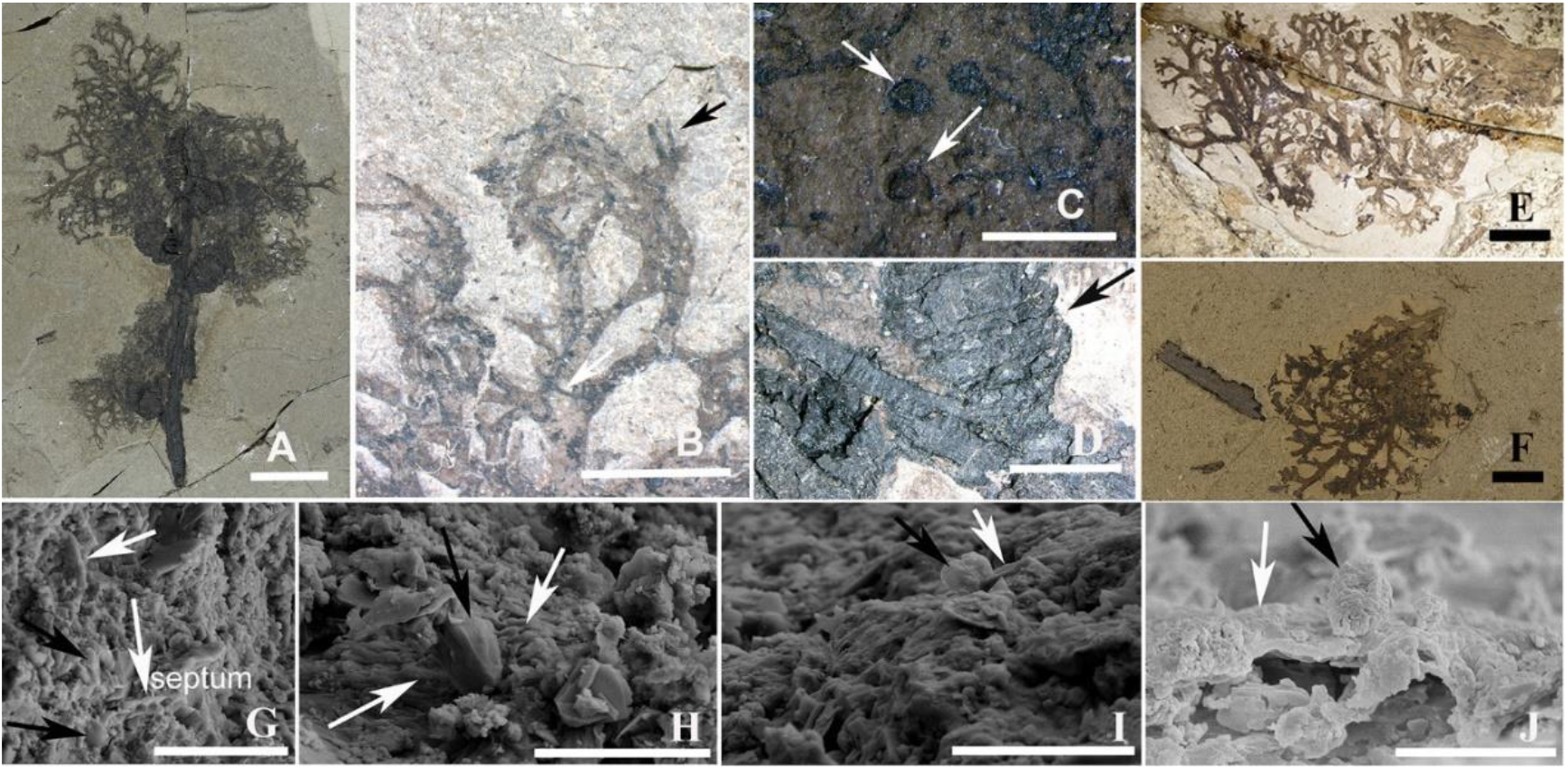
Photos of lichen *Daohugouthallus ciliiferus*. **(A)**. External morphology of lichen thallus directly growing on the gymnosperm branch, CNU-LICHEN-NN2020001. **(B)**. Marginal rhizinate cilia and lobules marked by white and black arrows, respectively, CNU-LICHEN-NN2020001; **(C)**. Superficial and nearly immersed unknown disc-like structure, CNU-LICHEN-NN2020001. **(D)**. Gymnosperm branch with seed cones marked by black arrow, CNU-LICHEN-NN2020001. **(E)**. External morphology of lichen thallus of CNU-LICHEN-NN2019001. **(F)**. External morphology of lichen thallus of CNU-LICHEN-NN2019002. Scanning electron microscopy (SEM) of *Daohugouthallus ciliiferus*. **(G–H)**. CNU-LICHEN-NN2020001. (**I)**. CNU-LICHEN-NN2020002. **(J)**. CNU-LICHEN-NN2019001. Scale bars: The fungal hyphae and algal cells indicated by white and black arrows, respectively. A, B, E=1cm; C, F=5mm; D=1mm; G=5 μm; H-J= 4 μm.

——Fungal Names FN570853

#### Diagnosis

Thallus corticolous, foliose to subfruticose, lobes irregularly branching, lateral black cilia and lobules present. Fungal hyphae thin, algal cells globose, one celled.

#### Type genus: Daohugouthallus

Wang, Krings & Taylor (*Wang et al., 2010*)

#### Type species: Daohugouthallus ciliiferus

Wang, Krings &Taylor (*Wang et al., 2010*)

Thallus foliose to subfruticose, about 5 cm high, 3 cm wide (*Fig. 1A, E, F*); lobes slender, about 5 mm long and 0.5–1.5 mm wide, tips tapering, nearly dichotomous to irregular branching, with lateral rhizinate cilia, concolorous to thallus to black, 0.5–1.5 mm long (*Fig. 1B*); surface yellow-brown with black spots in some area; lobules present (*Fig. 1B*); unknown disc-like structure superficial, or nearly terminal, yellow-brown, 0.25–0.5 mm in diam., sometimes immersed (*Fig. 1C*). Upper cortex conglutinate, comprising one cell layer, very thin, c. 1 μm thick (*Fang et al., 2020*); algal cells globose to near globose, one-celled, mostly 1.5–2.1 μm in diameter, some in framboidal form, anastomosed by or adhered to the fungal hyphae with simple wall-to-wall mycobiont-photobiont interface; fungal hyphae filamentous, some shriveled, septate, 1.2–1.5 μm wide (*Fig. 1G–J*; *Fang et al., 2020*).

#### Substrate

An unidentified gymnosperm branch (*Fig. 1D*).

#### Specimens examined

China, Inner Mongolia, Ningcheng County, Shantou Township, near Daohugou Village, Daohugou 1, Jiulongshan Formation, Callovian–Oxfordian boundary interval, latest Middle Jurassic. CNU-LICHEN-NN2019001, CNU-LICHEN-NN2020001, CNU-LICHEN-NN2020002, B0476P.

#### Remarks

The genus *Daohugouthallus* and species *D. ciliiferus* were published almost a decade ago (*Wang et al., 2010*); however, new anatomical characters were described only recently, based on scanning electron microscopy (*Fang et al., 2020*). Based on additional materials, phenotypic characters have been reassessed, including morphology and anatomy, and new information about the substrate ecology of the fossil has been added. Structural outline analysis of the thallus included not only the new material of the fossil but also a broad sampling of macrolichens, i.e. 61 extant species and two additional fossils from 12 families and six orders of Lecanoromycetes (*Table S1*). The geometric morphometric analysis of 140 images resulted in cumulative values for all the principal components were listed in *Table S2*. The cumulative eigenvalues for the main axes (principal components) with the cumulative variance of the first four principal components amounting to 61.1% (*Table S2*), meeting the requirements for geometric morphometric analysis. Among the canonical variate analysis (CVA) for four combinations of the four principal components (individual variances 24.4, 18.0, 10.8,8.0; *Fig. 2*), the plot combining the first two principal components (cumulative variance 42.4) showed that the fossil *Daohugouthallus ciliiferus*, in group 2, appeared morphologically closest to foliose Parmeliaceae, in group 3, including the genera *Hypotrachyna* and *Hypogymnia* and two foliose Parmeliaceae fossils (*Kaasalainen et al., 2017; Lücking and Nelsen, 2018*). However, placement of the fossil within the family Parmeliaceae is not possible, as the inferred stem age of Parmeliaceae is much younger than the age of the fossil (*Kraichak et al., 2018; Nelsen et al., 2020*). Therefore, a new family, Daohugouthallaceae is proposed, which is tentatively placed in the order Lecanorales given the close morphological similarity to Parmeliaceae.

**Fig. 2.**
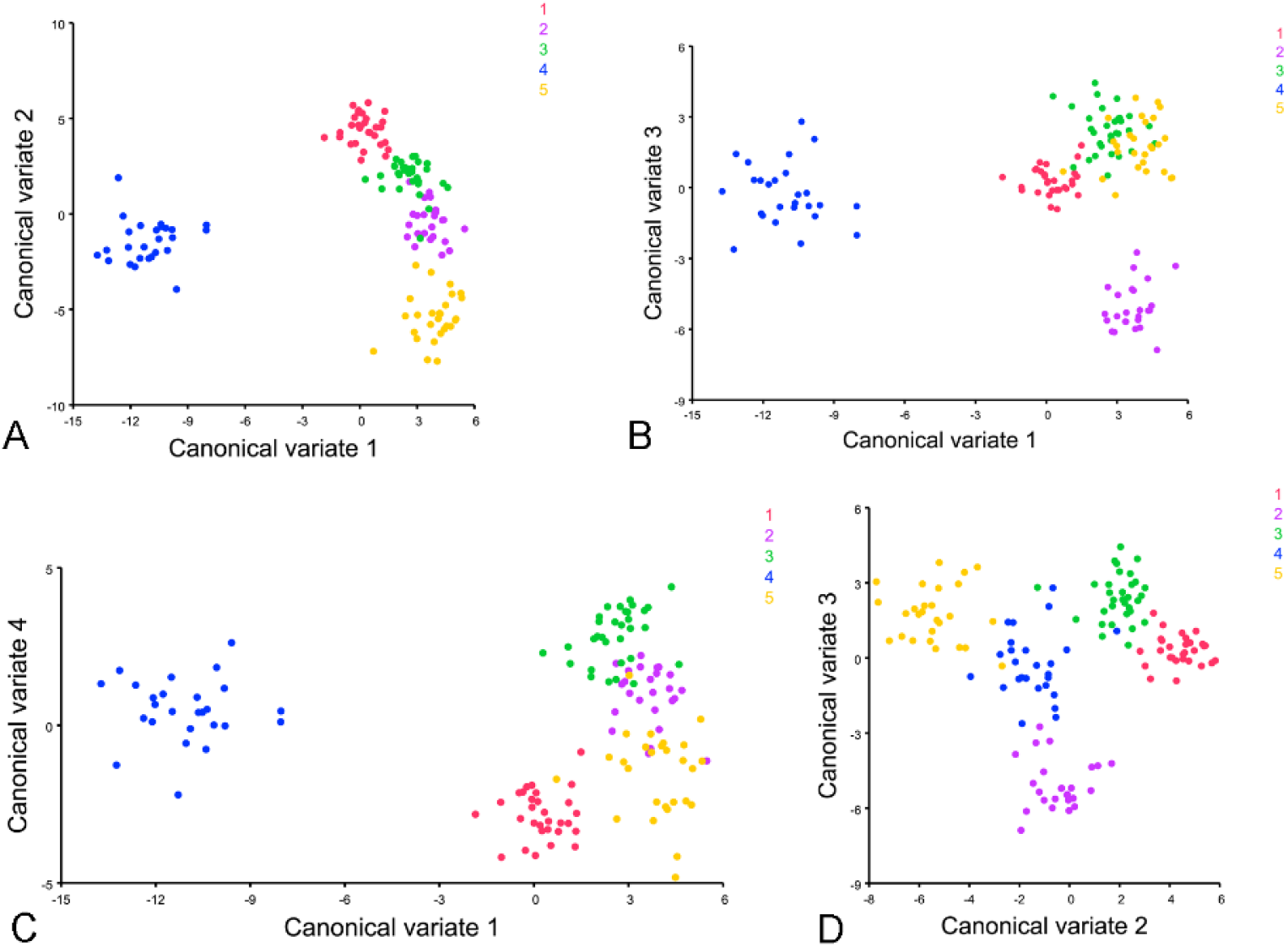
CVA plots based on the geometric morphometrics analysis (the first four principal components). **(A)**. CVA plot based on the highest Cumulative value 42.385 corresponding to the sum of the first principal component with the Variance value 24.433 and the second (17.952). **(B)**. CVA plot based on the Cumulative value 35.271 corresponding to the sum of the first principal component with the Variance value 24.433 and the third (10.838). **(C)**. CVA plot based on the Cumulative value 32.457 corresponding to the sum of the first principal component with the Variance value 24.433 and the fourth (8.024). **(D)**. CVA plot based on the Cumulative value 28.79 corresponding to the sum of the second principal component with the Variance value 17.952 and the third (10.838). Different colors represented different groups (*Table S3*). The distance showed the degree of similarity between different groups. Group 1 in red: FO-C-*Physcia* (Physciaceae, Caliciales)/FO-P-*Coccocarpia* (Coccocarpiaceae, Peltigerales)/FO-P-*Pannaria* (Pannariaceae, Peltigerales), group 2 in purple: FO-L-*Daohugouthallus*, group 3 in green: FO-L-Accepted foliose Parmeliaceae fossil/*Hypotrachyna*/*Hypogymnia* (Parmeliaceae, Lecanorales), group 4 in blue: FO-P-*Peltigera* (Peltigeraceae,Peltigerales)/*Lobaria*/*Sticta* (Lobariaceae, Peltigerales)/FO-U-*Umbilicaria* (Umbilicariaceae, Umbilicariales), group 5 in orange: FR-L-Accepted fruticose Parmeliaceae fossil/*Cladonia* (Cladoniaceae, Lecanorales) /*Evernia* (Parmeliaceae, Lecanorales)/*Ramalina* (Ramalinaceae, Lecanorales)/*Sphaerophorus* (Sphaerophoraceae, Lecanorales)/FR-P-*Siphula* (Icmadophilaceae, Pertusariales)/FR-T-*Teloschistes* (Teloschistaceae, Teloschistales).

### Significance of the occurrence of Daohugouthallaceae

We used the detailed molecular clock tree provided by *Nelsen et al. (2020)* to illustrate inferred ages for selected family-level clades in the Lecanoromycetes that include macrolichens (*Fig. 3*). Most of these families have stem node ages relative to the macrolichen genera contained therein substantially younger than 100 Mya, including Caliciaceae, Cladoniaceae, Pannariaceae, Parmeliaceae, Physciaceae, Ramalinaceae, Sphaerophoraceae, Stereocaulaceae and Teloschistaceae. The stem node ages of a few families including macrolichens were reconstructed as between 150 and 100 Mya, including Baeomycetaceae, Coccocarpiaceae, Collemataceae, Pannariaceae, Peltigeraceae, and Umbilicariaceae. However, all these families have a crown node age significantly younger than 165 Mya, the age of Daohugouthallaceae (*Fig. 4*). The only macrolichen family older than the fossil is Icmadophilaceae, with an inferred crown node age of approximately 200 Mya (*Figs 3–4*). However, members of this family differ strongly in ecology and morphology from Daohugouthallaceae (*Fig. 4*). In addition, the macrolichen genera within Icmadophilaceae distinctly diversified after the K–Pg boundary: *Siphula* approximately 48 Mya and *Thamnolia* about 16 Mya (*Fig. S1*). If these estimates are correct, the Jurassic macrolichen cannot be included in any extant family containing macrolichens, and so the occurrence of Daohugouthallaceae in the Jurassic may reflect a scenario of early macrolichen diversification long before the diversification of extant lineages of epiphytic macrolichens.

**Fig. 3.**
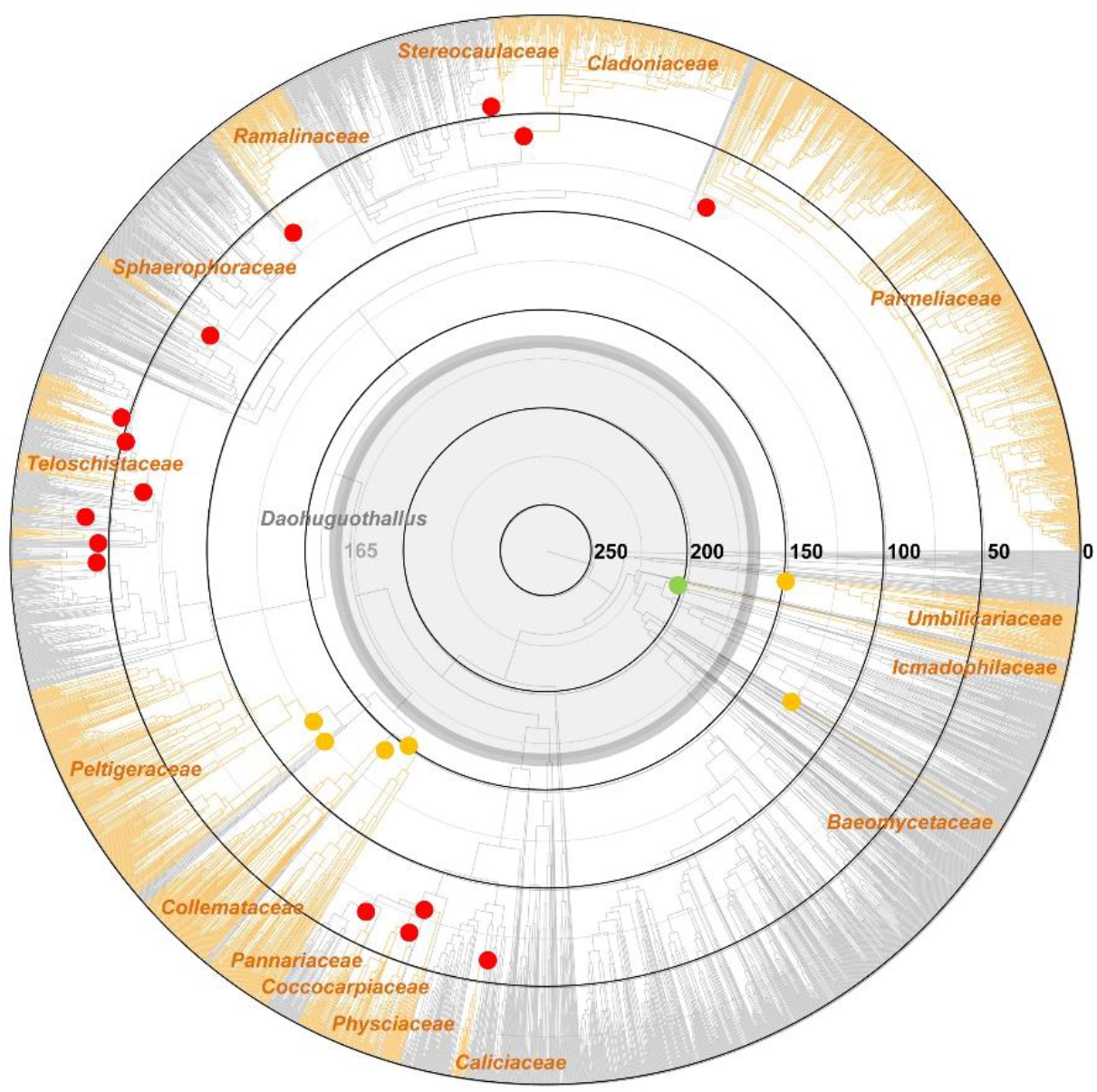
Time-calibrated ML phylogeny of 3,373 Lecanoromycetes fungi (based on *Nelsen et al., 2020*). Macrolichen lineages are indicated in orange and the corresponding families are indicated. The temporal placement of the Daohugouthallus ciliiferus fossil is marked by the bold gray circle. For details see Suppl. Fig. S1.

**Fig. 4.**
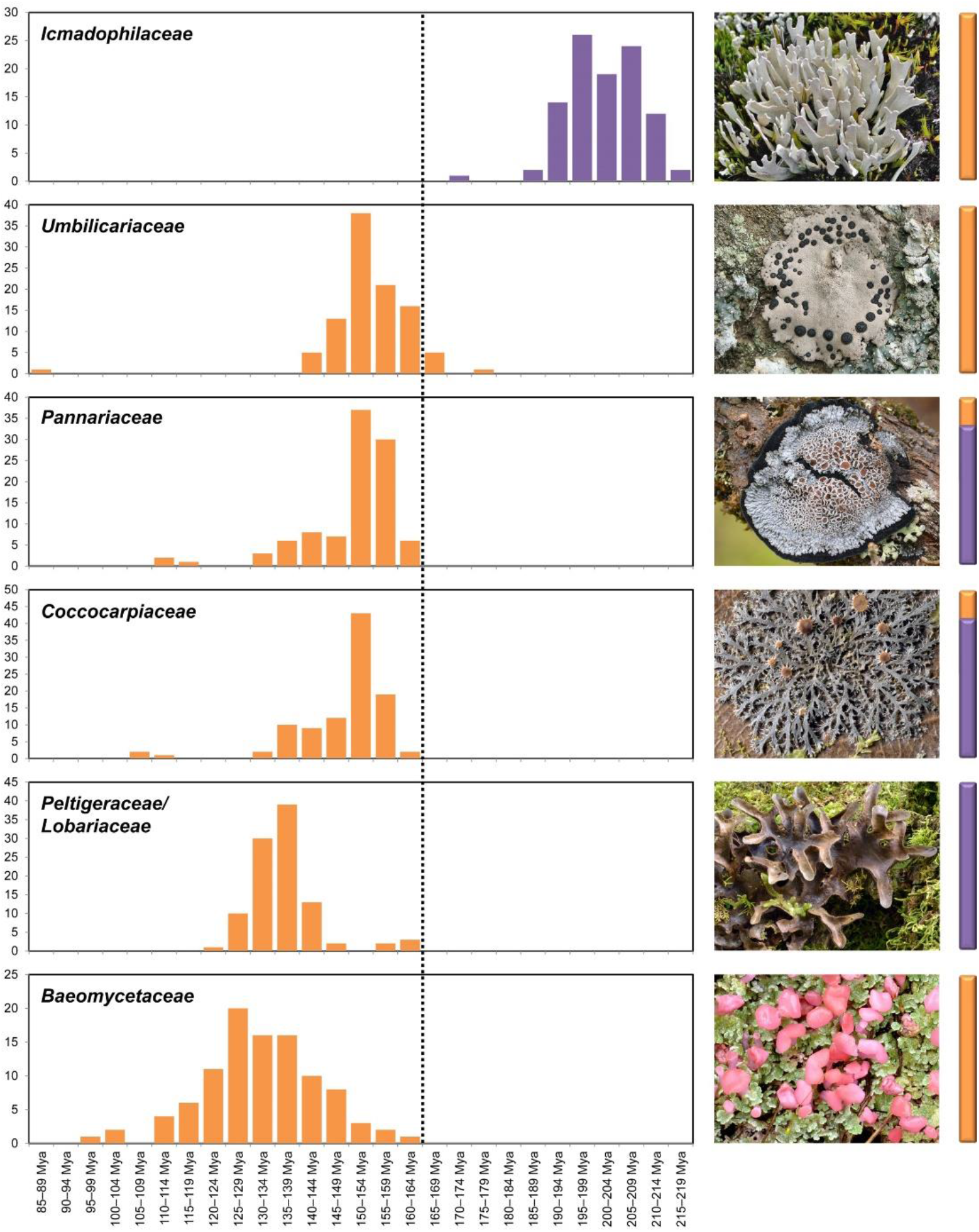
Distribution of inferred divergence times for the oldest extant macrolichen families (based on *Nelsen et al., 2020*). The external morphology of selected representatives of each family is depicted to the right. The dotted line indicates the temporal placement of the *Daohugouthallus ciliiferus* fossil. The first three families are likely as old or older than the fossil but do not fit morphologically and/or ecologically. The best morphological and ecological fit are Peltigeraceae, in particular lobarioid lineages, but that family is significantly younger.

**Fig. 5.**
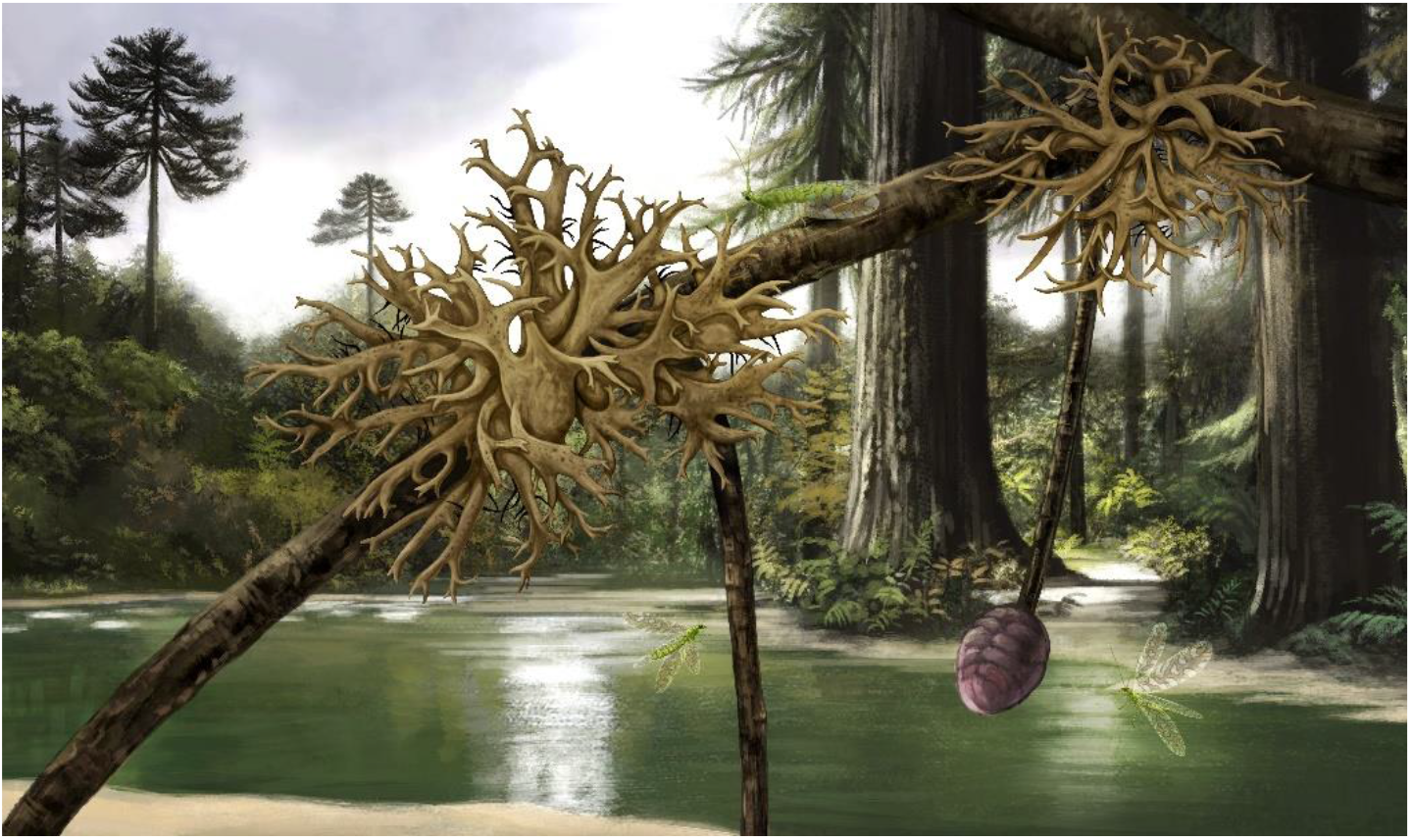
Habitus reconstruction of the lichen *Daohugouthallus ciliiferus* growing on gymnosperm branches. Drawing by Xiaoran Zuo.

## Discussion

### The phylogenetic placement of Daohugouthallaceae

The new family based on the fossil lichen *Daohugouthallus ciliiferus* is tentatively placed in Lecanorales, although sexual reproductive structures are missing that would allow to test this placement. Ascomata, asci and ascospores are crucial to assess systematic affinities within Lecanoromycetes (*Hafellner, 1994*) and have been demonstrated in fungal fossils as old as 400 Mya (*Taylor et al., 2005*). Apothecia-like structures seem to be present in the fossil (*Fig. 1C*), but no structures interpretable as asci and ascospores were detectable. The thin nature of compression fossils like *Daohugouthallus ciliiferus* makes it difficult for such structures to be preserved, if they were indeed present in the organism. Molecular phylogeny has also shaped the systematics of Lecanoromycetes (*Thell et al., 2009; Miadlikowska et al., 2011*; *Leavitt et al., 2015; Magain et al., 2017*). DNA has been presumably extracted and amplified from fossils as old as 250 Mya (*Cano et al., 1993; Fish et al., 2002*), but these findings have been challenged and considered artifactual (*Pääbo et al., 2004*). Successful DNA extraction from permineralized or compression fossils as old as *Daohugouthallus ciliiferus* seems impossible and so this is not an avenue that could be followed to clarify the systematic affinities of this and other lichen or fungal fossils.

In lieu of sexual reproductive structure evidence to clarify the potential affinities of Daohugouthallaceae, our geometric morphometric analysis seems a suitable alternative to provide at least a hypothesis based on quantitative data. This approach, based on homologous landmarks or structural outlines (*Rohlf and Marcus, 1993*), was here apparently used for the first time in this context but has been widely used in entomology (*Bai et al., 2012, 2014; Ren et al., 2017*). However, it requires a careful approach to data assessment (*Fox et al., 2020*). The CVA plots (*Fig. 2*) based on the comparison with homologous landmarks of 61 extant macrolichens and two Parmeliaceae fossils showed that Daohugouthallaceae are most similar to foliose Parmeliaceae (Lecanorales). Given the much younger stem age of the latter (and other related families such as Physciaceae), the introduction of a new, monogeneric family for this fossil therefore seems justified.

The apparently smaller algal cells (1.5–2.1 μm in diam.) and thinner fungal hyphae (1.2–1.5 μm wide) in *Daohugouthallus ciliiferus* (*Fig. 1G–J*; *Fang et al., 2020*) compared to extant Lecanoromycetes, suggest that Daohugouthallaceae may have also originated from a relict clade outside Lecanoromycetes; however, it is unclear how the fossilization process may have affected these structures.

### The importance of Daohugouthallaceae for our understanding of macrolichen evolution

Among the new specimens of *Daohugouthallus ciliiferus* collected from the type locality, one thallus was found attached to the branch of an unidentified cone-bearing gymnosperm fossil. The Daohugou paleoenvironment has been analyzed to be a gymnosperm-dominated forest vegetation (*Ren and Krzeminski, 2002; Tan and Ren, 2002; Zhang et al., 2006*), and *Daohugouthallus ciliiferus* has been reconstructed to be epiphytic on gymnosperms due to its association with a small seed cone (*Wang et al., 2010*), but in the original study the two fossils were not directly connected. Our new specimen (*Fig. 1A*) clearly shows the thallus growing directly on a thin branch of a gymnosperm with an associated cone (*Fig. 1D*), possibly representing a conifer, suggesting that gymnosperms may have served as substrate for epiphytic macrolichens already in the Jurassic. Extant lichenized clades that are largely epiphytic have been dated back to as far as the early Jurassic, such as microlichens in the Graphidaceae, Pyrenulaceae, and Trypetheliaceae (*Lücking et al., 2013; Beimforde et al., 2014; Kraichak et al., 2018; Nelsen et al., 2019, 2020*). However, the oldest extant macrolichen clade, *Umbilicaria*, is almost exclusively rock-dwelling (saxicolous), and while extant macrolichen vary greatly in substrate choice (*Belnap et al., 2001; Brodo et al., 2001; Nash, 2008*), extant epiphytic macrolichen clades are consistently younger than the K–Pg boundary and only since then have evolved to form the conspicuous elements of terrestrial woody ecosystems they are today (*Ahti et al., 1993; Esslinger,1985; Goward and McCune, 2007; McCune et al., 2000; Spribille and Muggia, 2013; Wei et al., 2017*). The fossil record and molecular clock studies indicate that gymnosperms diverged around 315 Mya (*Crisp and Cook, 2011*; *Nie et al., 2020*), while conifers originated approximately 300 Mya and diversified between 190–160 Mya in the Early to Middle Jurassic (*Leslie et al., 2018*) into the various families recognized today. The divergent time of Lecanoromycetes, the main class including epiphytic macrolichens, has been estimated at 300–250 Mya (*Nelsen and Lücking, 2018; Kraichak et al., 2018; Lutzoni et al., 2018*; *Nelsen et al., 2020*), and so it is conceivable that epiphytic macrolichens even older than *Daohugouthallus ciliiferus* may have existed.

The diversification of Lecanoromycetes coincides with the period after the end-Permian extinction. Before that time, diverse Permian forests existed around the world (*Behrensmeyer and Hook, 1992; Gulbranson et al., 2012; Wang et al., 2012*). While these could have provided potential environments for epiphytic macrolichens, there is no fossil record that would support such an assumption. The end-Triassic mass extinction 200 Mya greatly affected marine and terrestrial organisms (*Erwin, 2001; Damborenea et al., 2017*), but its effect on lichenized fungi is unclear. The ecology of terrestrial vegetation at that time, with diverse forests already in the Late Triassic and into the Jurassic (*Rees et al., 2004; Wang et al., 2005; Sellwood & Valdes, 2008; Bonis & Kürschner, 2012*), would certainly have supported the existence of epiphytic macrolichens, but again, no fossil record exists that would support such as a hypothesis. In contrast, the diversification of the crown clades of extant conifers, between 190 and 160 Mya in the Early to Middle Jurassic (*Leslie et al., 2018*), coincides well with the 165 Mya *Daohugouthallus ciliiferus* fossil, thus providing the earliest known evidence of the existence of epiphytic macrolichens. Notably, various other groups of organisms underwent radiations in this period, such as mammals or the avian stem lineage (*Benson et al., 2014; Close et al., 2015*). The Mid-Mesozoic era was a cooling and greenhouse period (*Willis and Niklas, 2004*) and the palaeoenvironment of the Daohugou formation has been described as humid, warm-temperate and montane (*Ren and Krzeminski, 2002; Tan and Ren, 2002; Zhang et al., 2006*), thus favoring the potential growth of epiphytic macrolichens, as shown by the ecology of extant macrolichen lineages in the tropics. The presence of a fossil macrolichen such as *Daohugouthallus ciliiferus* is therefore not surprising and we expect that other macrolichen morphotypes may be found in this and other formations of the Mesozoic, hopefully expanding the evidence for the diversification of macrolichen lineage well before the most recent mass extinction event, the K–Pg boundary.

## Materials and Methods

### Geological context

Specimens in this study were collected from the Daohugou locality of the Jiulongshan Formation, near Daohugou Village, Ningcheng County, approximately 80 km south of Chifeng City, in the Inner Mongolia Autonomous Region, China (119°14.318′E, 41°18.979′N). The age of this formation is 168–152 Ma based on ^40^Ar/^39^Ar and ^206^Pb/^238^U isotopic analyses (*He et al., 2004; Liu et al., 2006; Ren et al., 2019*).

### Experimental methods

The lichen fossils were examined and photographed using an Olympus SZX7 Stereomicroscope attached to a Mshot MD50 digital camera system. For selected fossil we made cross sections using a stonecutter, then sputter-coated these with gold particles using Ion Sputter E-1045 (HITACHI). SEM images were recorded using a scanning electron microscope (Hitachi SU8010) with a secondary electron detector operated at 5.0 kV. Plates were composed in Adobe Photoshop. All lab work was performed at the Institute of Microbiology, Chinese Academy of Sciences in Beijing, except the stonecutter which was located at the Institute of Geology and Geophyscis, Chinese Academy of Sciences in Beijing.

### Specimen repository

Fossil specimens of *Daohugouthallus ciliiferus* (CNU-LICHEN-NN2019001, CNU-LICHEN-NN2020001, CNU-LICHEN-NN2020002, B0476P) are deposited in the Key Lab of Insect Evolution and Environmental Changes, College of Life Sciences and Academy for Multidisciplinary Studies, Capital Normal University, in Beijing, China.

### Geometric morphometrics

Absence of diagnostic characters such as ascomata, asci and ascospores in the adpression fossil *Daohugouthallus ciliiferus* made it difficult to place into extant lineages. Therefore, we focused on thallus characters, mainly the topology of lobes or branches, to implement geometric morphometric analysis, based on a curve connected by tracing the points of the ends of lobes or branches. The starting point of the curve was selected as a point on the upper edge of the lobe or branch near the center or substrate, and after describing the outline of the whole lobe or branch, the end point returning to the lower edge near the starting point. For this purpose, 113 images of 61 representative extant macrolichen species were selected from 12 families and 6 orders of Lecanoromycetes (*Table S1*), including specimens deposited in HMAS-L, photos provided by Robert Lücking, and pictures downloaded from the CNALH (Consortium of North American Herbaria) Image Library https://lichenportal.org/cnalh/imagelib/ and the *Hypogymnia* Media Gallery http://hypogymnia.myspecies.info/gallery, together with two images of accepted Parmeliaceae fossils (*Kaasalainen et al., 2017*).

A total of 140 images (*Fig. S3*), including the above-mentioned images and the *Daohugouthallus ciliiferus* fossil images, were cut into 25 sub-images and used as test groups. The whole image set was divided into five groups according to lobes types: microfoliose group, the *Daohugouthallus ciliiferus* fossil group, a long branches group, a wide-lobed group, and a fruticose group (*Table S3*). The selected images were two-dimensional graphs with two views of the front or back of the thallus where the branch tips were clearly recognizable. To orientate the images in the same direction, they were adjusted so that the end of the branches faced right. Images were named in a unified format: growth type-order-family-genus-species (sample number) except for the two selected reference fossil images only corresponding to family name.

As stated, the external forms were represented by one curve extracted from the end of branches or lobes and the curve was resampled into 60 semi-landmarks by length (*Fig. S2*). The curves and semi-landmarks were digitized using TPS-DIG 2.05 (*Rohlf, 2006*). To merge all semi-landmarks into the same data file to produce the data set for morphological analysis, the data file was opened as text file to convert the semi-landmarks to landmarks, by deleting the line with the curve number and point number and replacing the landmark number by the point number (*Tong et al., 2021*).

MORPHO J 1.06a (*Klingenberg, 2011*) was used for subsequent analysis of the data set. Through Procrustes analysis, the morphological data of all test features were placed in the same dimensional vector space to screen out physical factors such as size. Principal component analysis (PCA) and geometric modeling of the mathematical space formed by PC axis were used to coordinate the shape changes of the entire dataset. We then selected the data set to generate a covariance matrix. In this context, the first two principal components corresponding to the highest cumulative variance represent the best variation pattern of test shape. The relationships among different morphological groups were then visualized through canonical variate analysis (CVA).

## General

We thank Mr. Yijie Tong for assisting in geometric morphometrics analysis, Dr. Chunli Li for assisting in taking SEM photos, Ms. Hong Deng and Ms. Haijuan Chen for lending and taking photos of lichen specimens in HMAS-L, Ms. Xiaoran Zuo for drawing the habitus reconstruction picture *Fig. 5*, and Mr. Jujie Guo for making the fossil slices.

## Funding

This work was supported by the National Natural Science Foundation of China (grant 31770022 and 32070096 to X.W.; grant 31970383 to Y.W.; grants 31730087, 41688103, 32020103006 to D.R.; grant 31961143002 to M.B.) and Ministry of Science and Technology of China (grant 2019FY101808 to X.W.); the Beijing Natural Science Foundation (grant 5192002 to Y.W.).

## Author contributions

QXY: Literatures investigation, geometric morphometric analysis, molecular data collection and analysis, original draft editing; YYW: SEM photos taking and analysis, molecular data analysis, original draft editing; RL: Conceptualization, molecular analysis, validation, draft review and editing; TL: Conceptualization, validation, draft review and editing; XW: Fossil lichen and plant identification, draft review and editing; ZYD: molecular analysis; YKC: Geometric morphometric analysis; MB: Geometric morphometric analysis, draft review and editing; JCW: Conceptualization, supervision, validation, draft review and editing; DR: Conceptualization, resources collection, supervision, project administration, validation, funding acquisition, draft review and editing; HL: Conceptualization, molecular analysis, validation, draft review and editing; YJW: Conceptualization, resources collection, supervision, funding acquisition, validation, draft review and editing; XLW: Conceptualization, supervision, geometric morphometric analysis, SEM photos taking and analysis, data curation, funding acquisition, project administration, validation, original draft writing, review and editing.

## Notes

### Competing Interest Statement

The authors have declared no competing interest.

