## Supplementary Materials for "New insights into the earlier evolutionary history of epiphytic macrolichens"

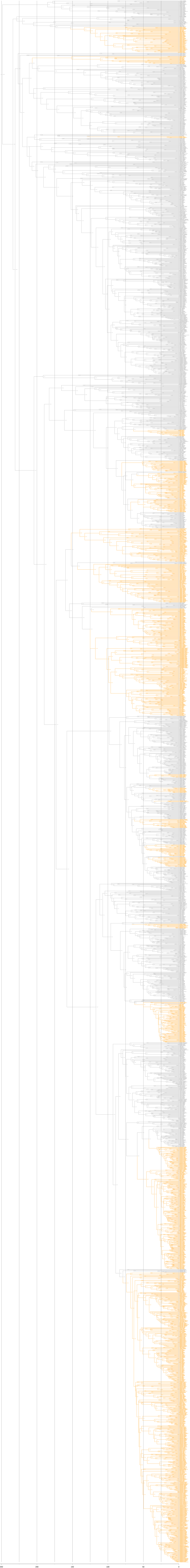
250
200
150
100
50
0

**Fig. S1.** Detailed time-calibrated ML phylogeny of 3,373 Lecanoromycetes fungi (based on *Nelsen et al., 2020*). Macrolichen lineages are indicated in orange.

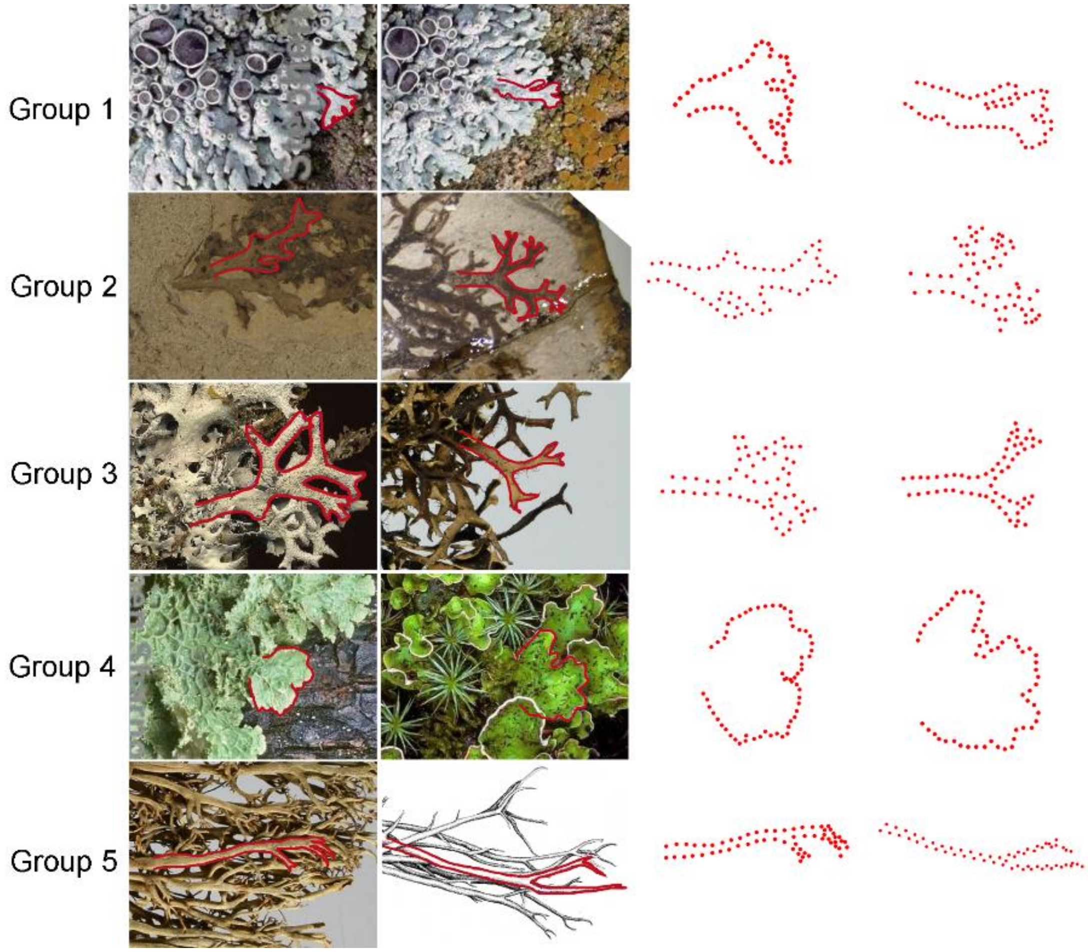

**Fig. S2**. Description of the curves used in the geometric morphometric analysis. The curves of lobes or branches of the five groups (small branches group, *Daohugouthallus ciliiferus* fossil group, long branches group, wide lobes foliose group and fruticose group) shown in columns 1 and 2 are outlined in red color, then they were resampled in 60 semi-landmarks represented by red points shown in columns 3 and 4.

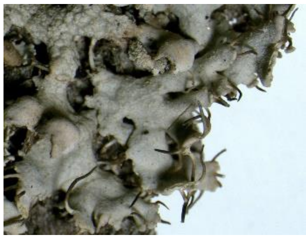
0

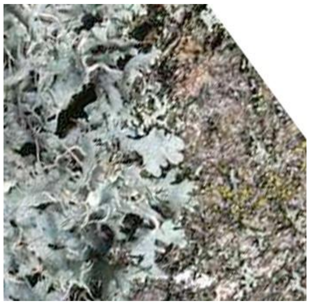
1

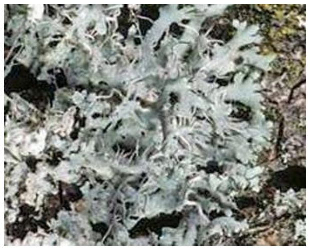
2

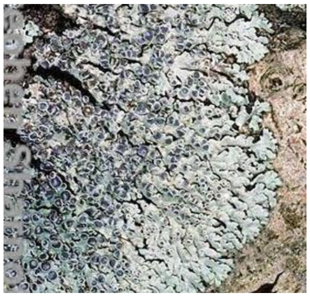
3

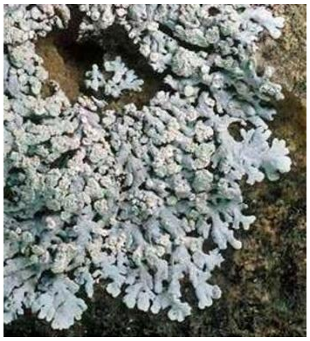
4

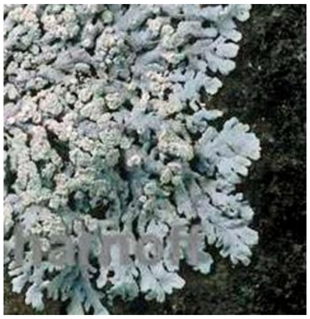
5

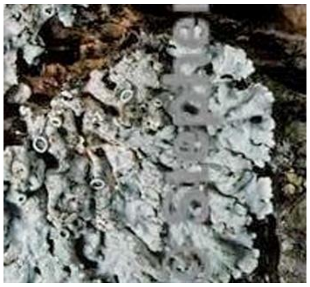
6

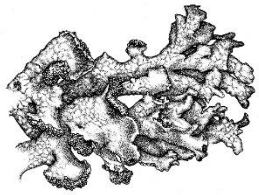
7

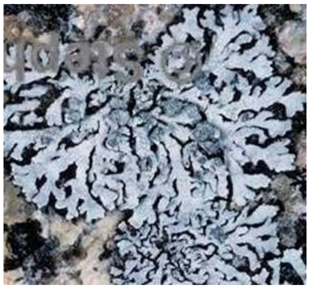
8

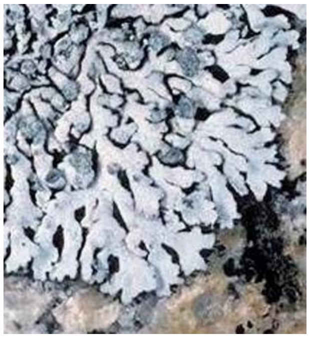
9

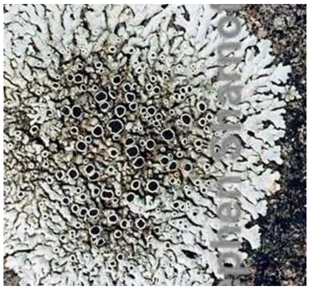
10

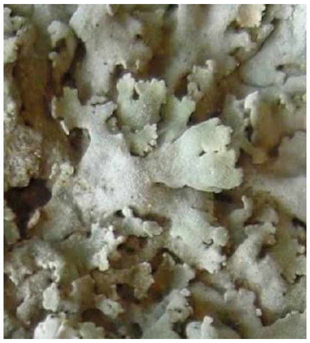
11

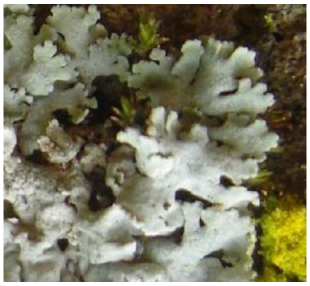
12

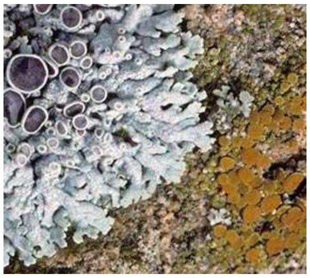
13

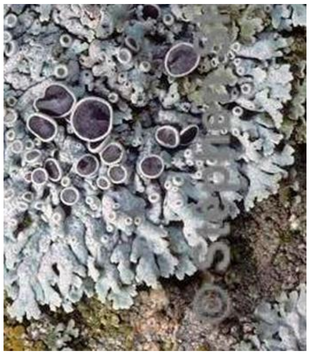
14

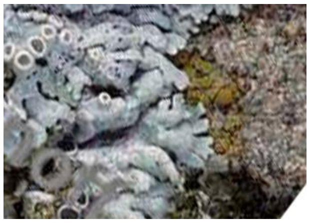
15

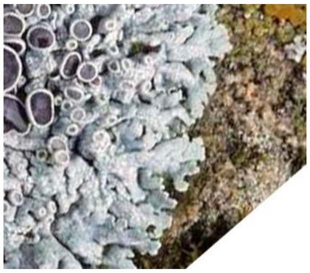
16

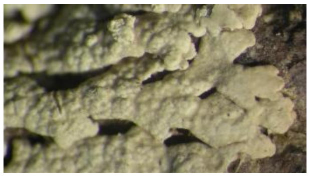
17

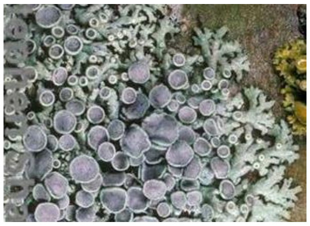
18

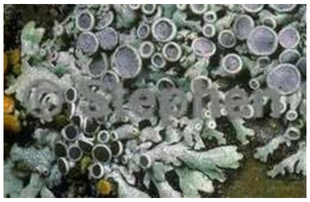
19

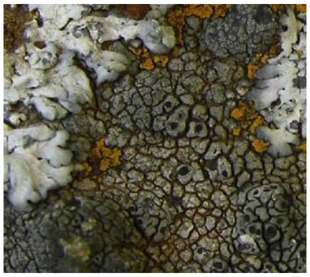
20

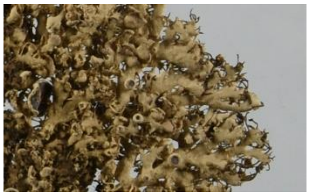
21

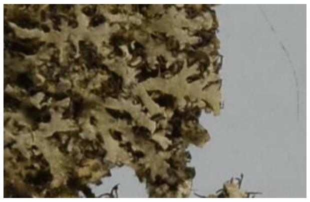
22

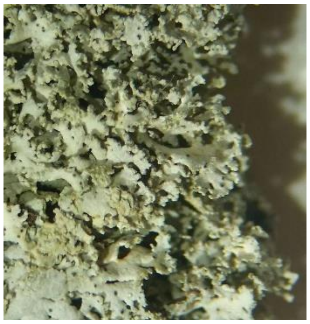
23

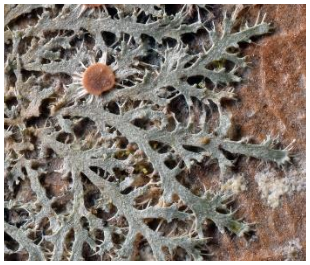
24

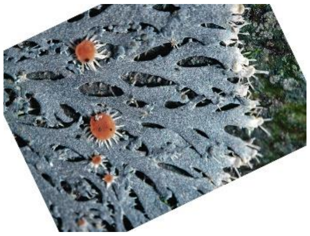
25

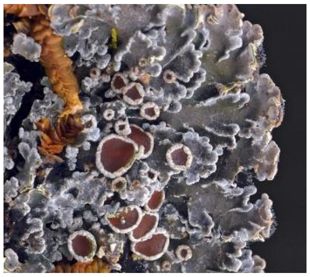
26

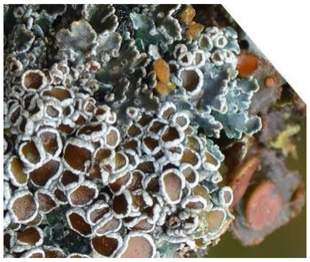
27

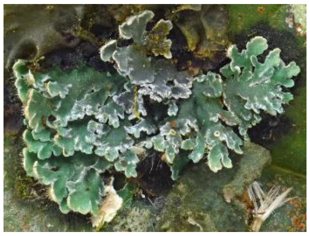
28

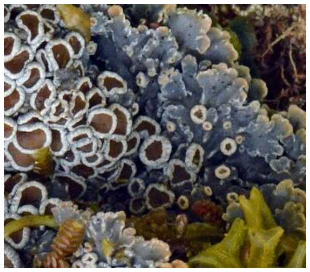
29

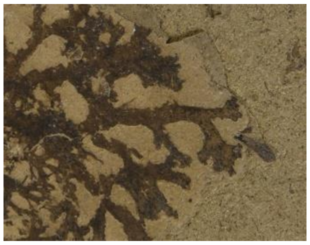
30

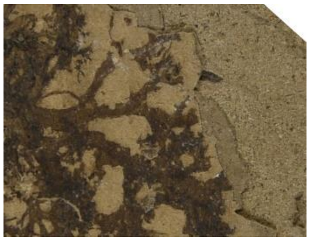
31

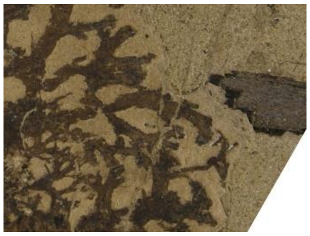
32

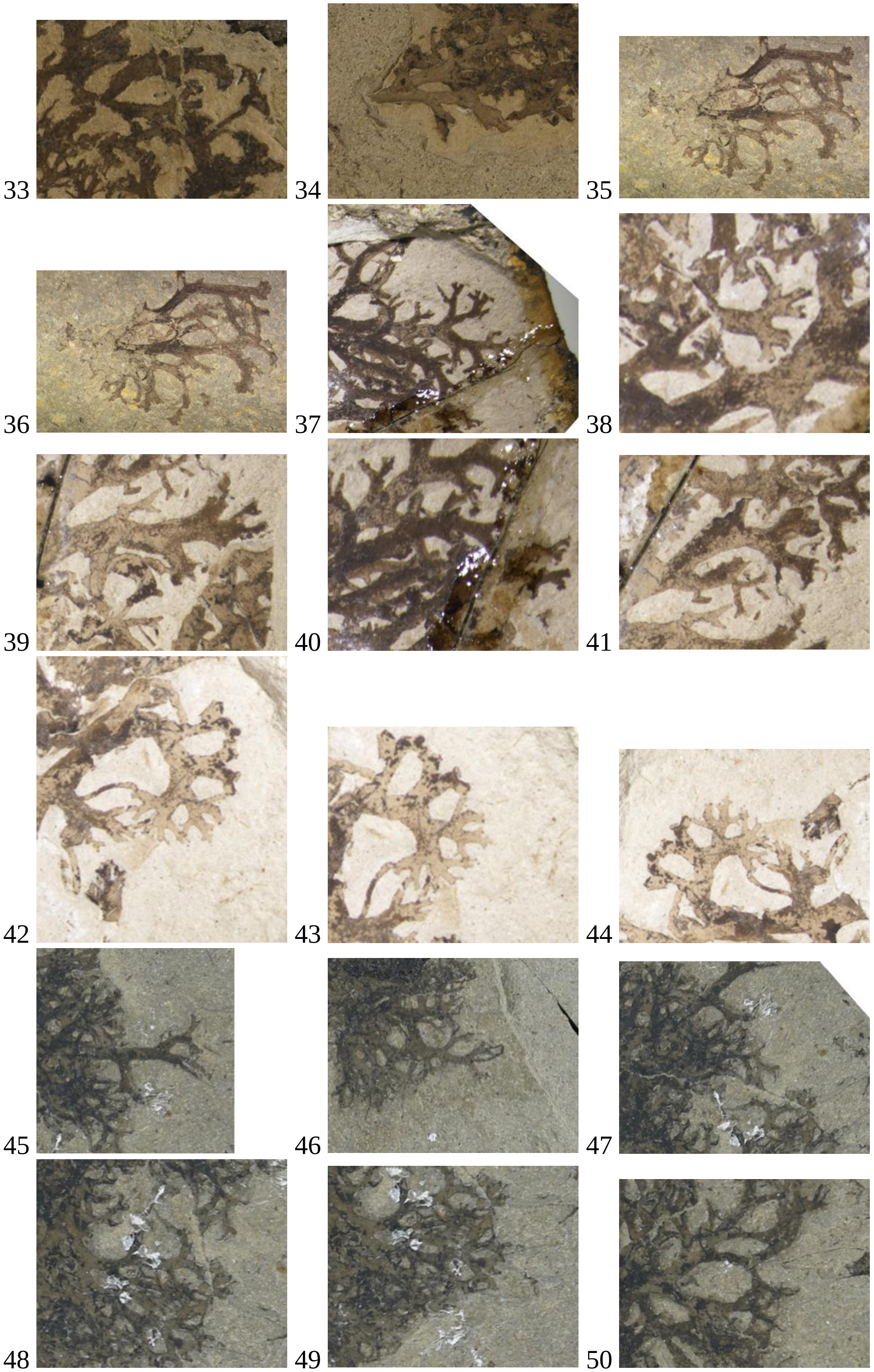

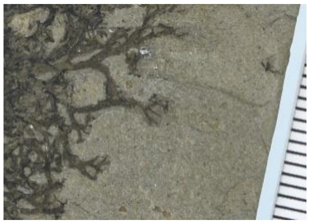
51
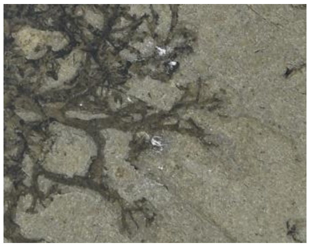
52
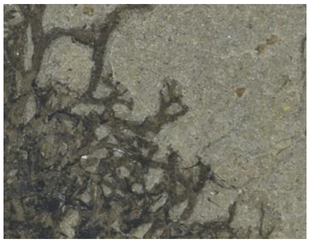
53
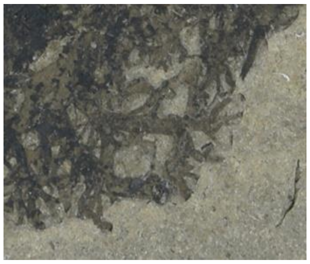
54
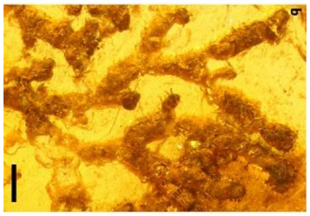
55
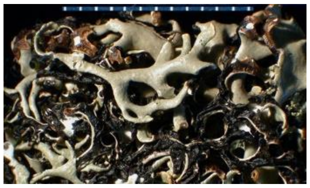
56
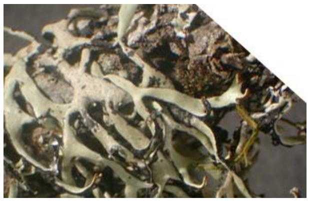
57
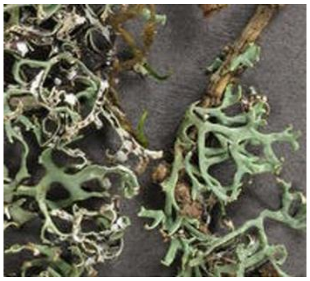
58
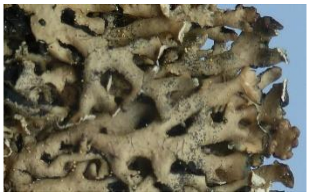
59
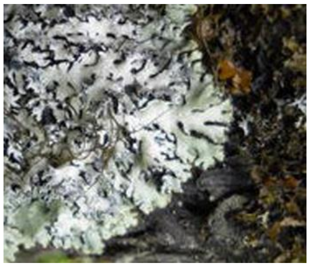
60
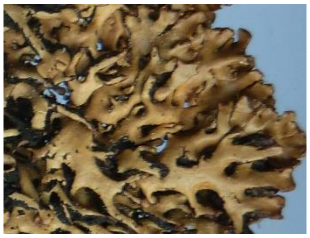
61
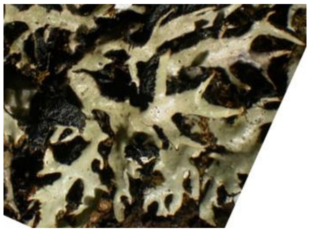
62
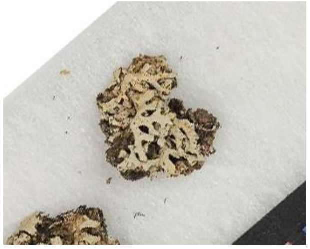
63
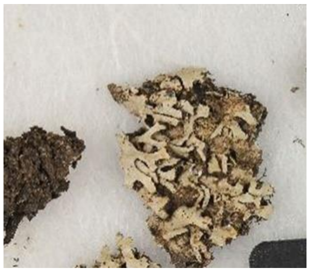
64
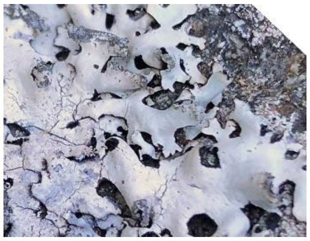
65
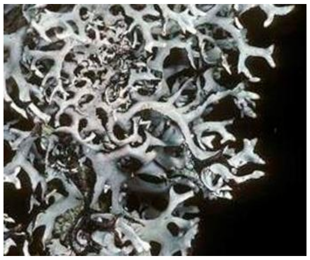
66
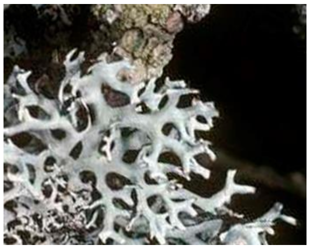
67
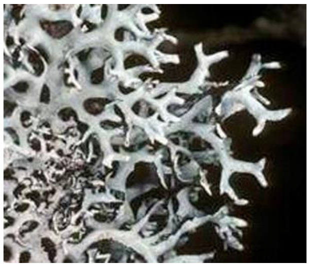
68

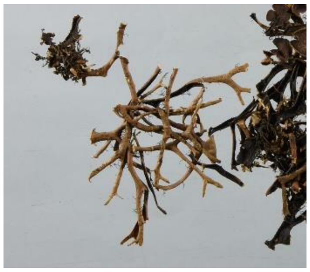
69

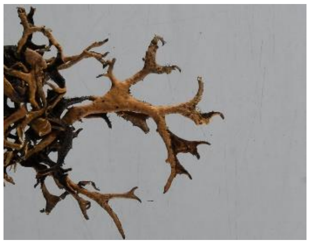
70

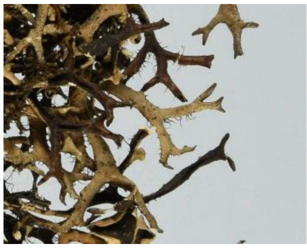
71

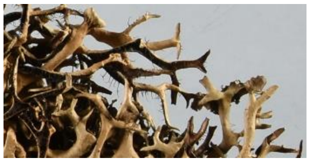
72

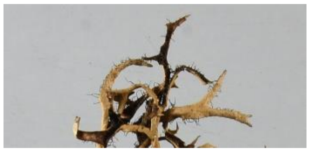
73

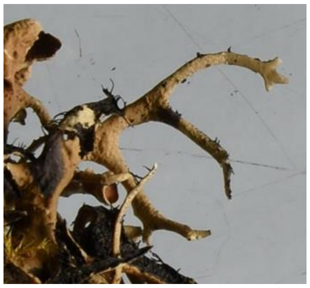
74

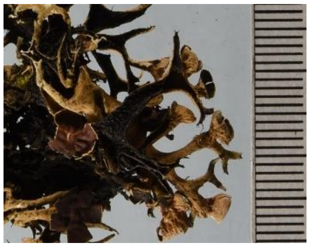
75

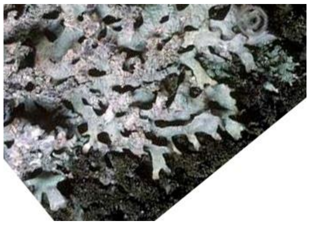
76

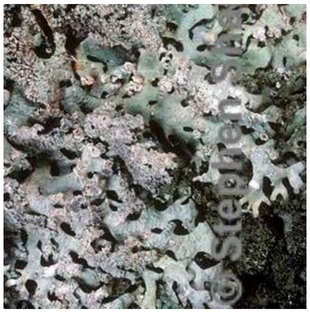
77

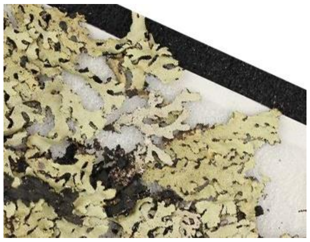
78

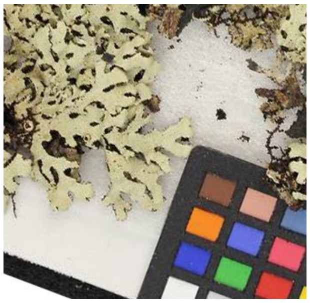
79

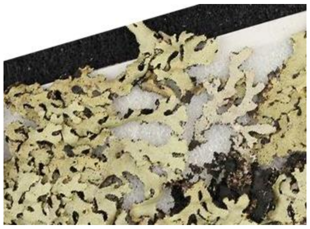
80

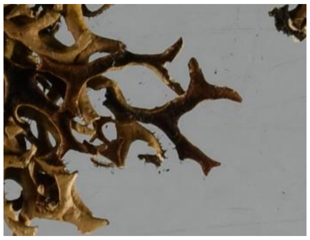
81

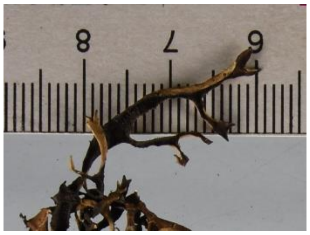
82

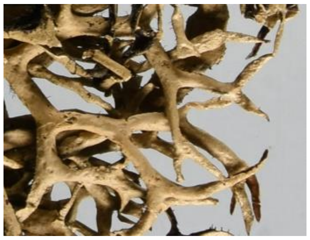
83

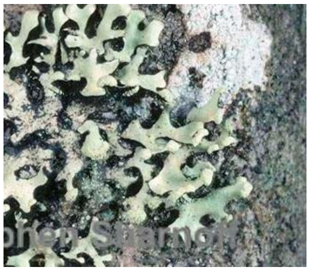
84

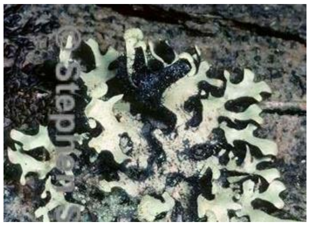
85

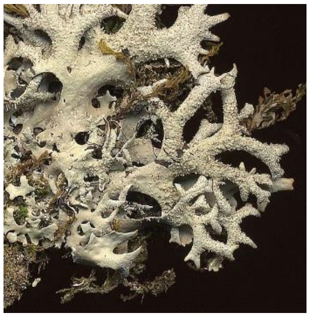
86

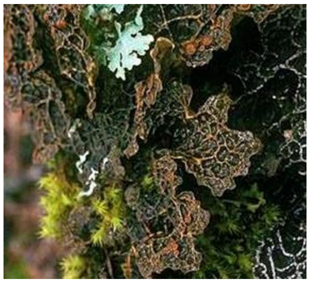
87

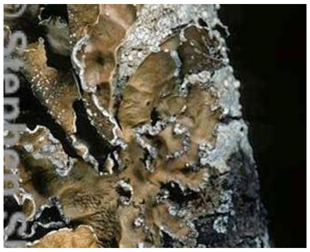
88

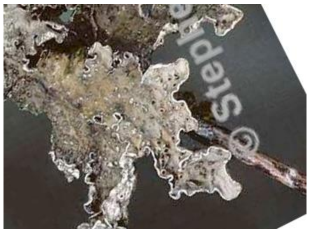
89

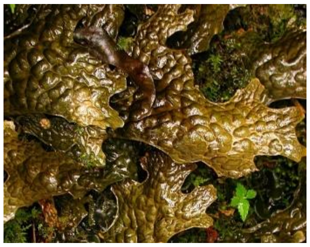
90

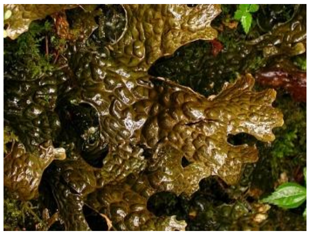
91

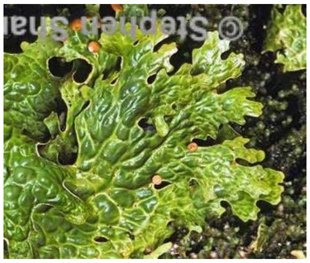
92

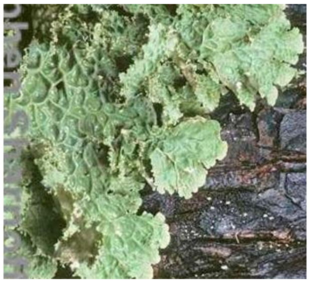
93

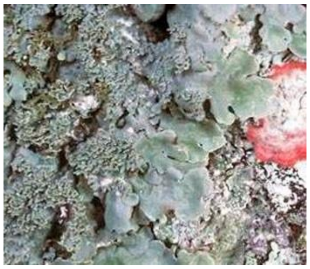
94

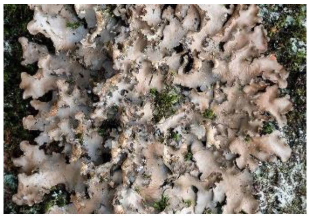
95

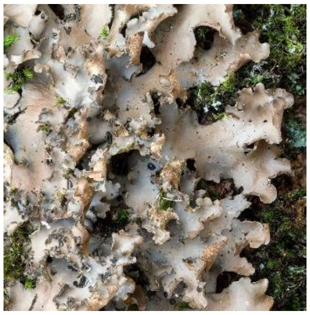
96

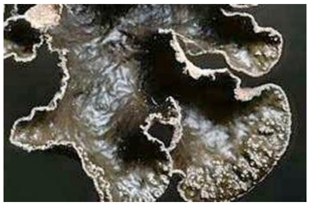
97

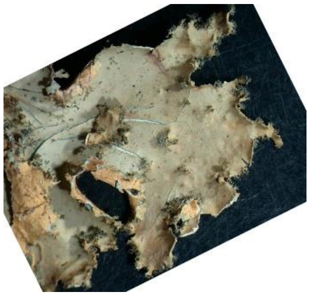
98

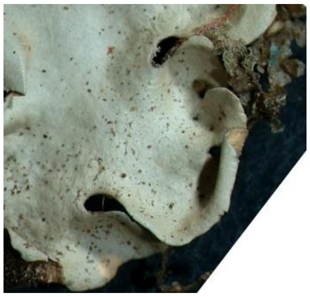
99

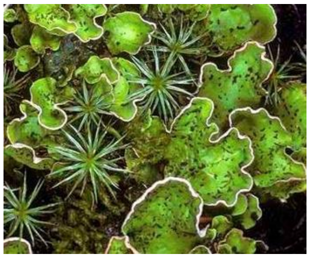
100

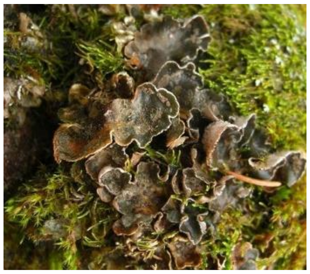
101

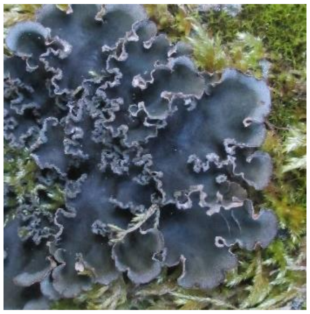
102

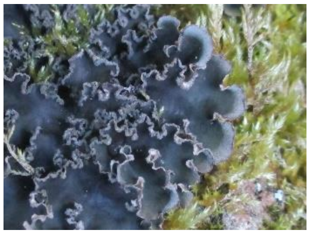
103

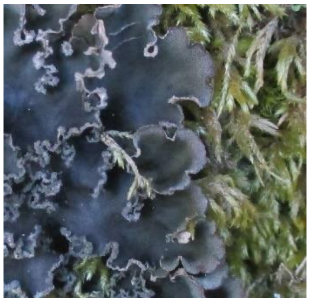
104

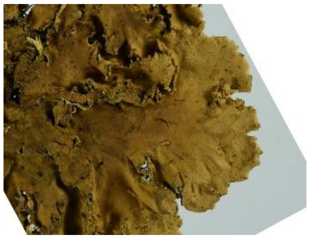
105

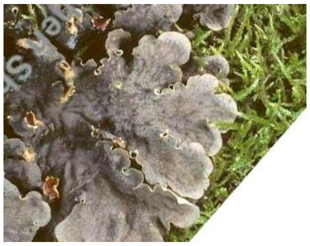
106

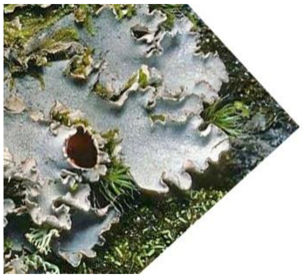
107

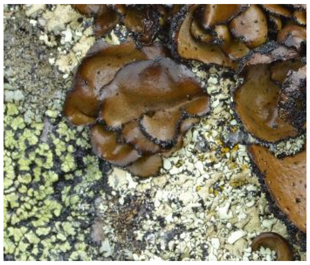
108

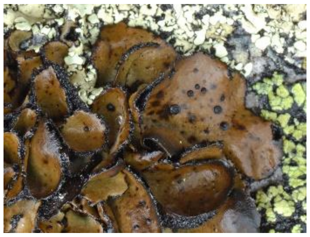
109

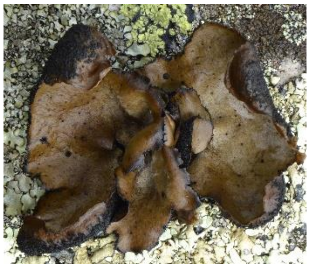
110

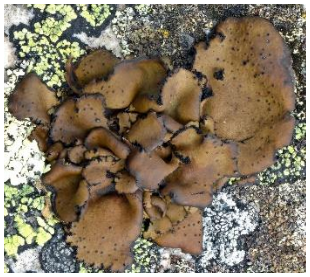
111

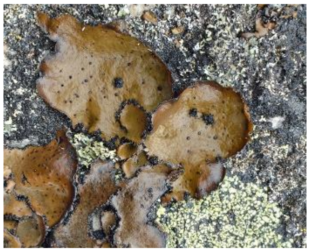
112

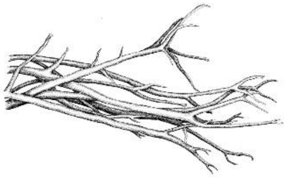
113

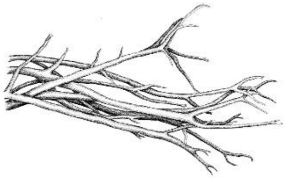
114

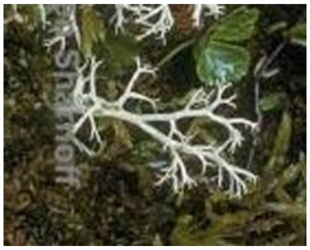
115

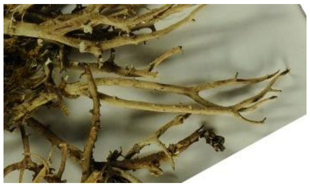
116

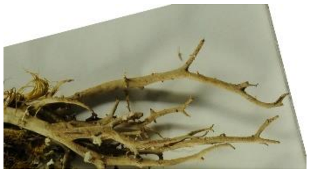
117

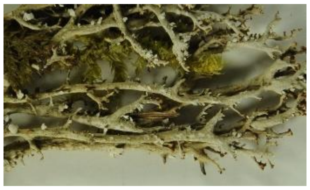
118

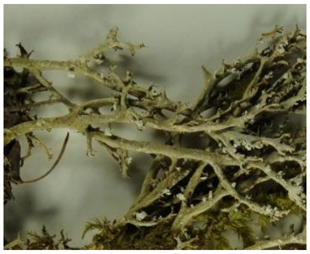
119

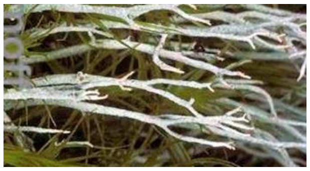
120

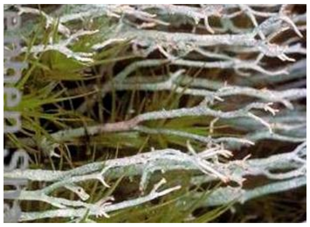
121

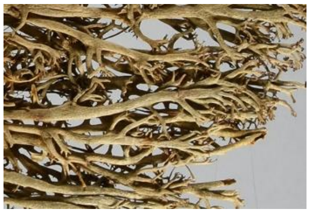
122

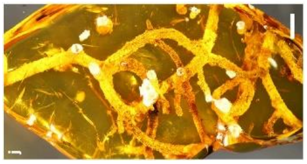
123

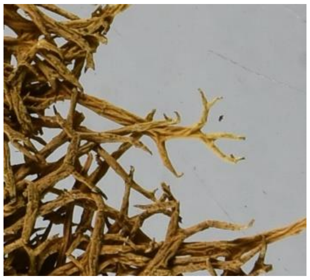
124

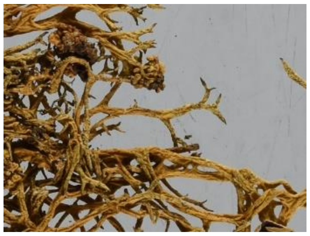
125

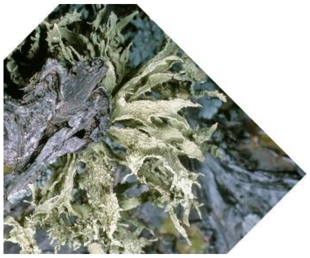
126

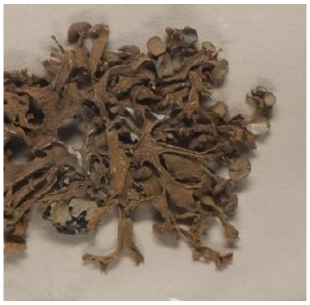
127

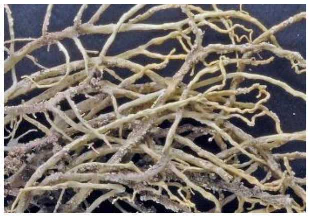
128

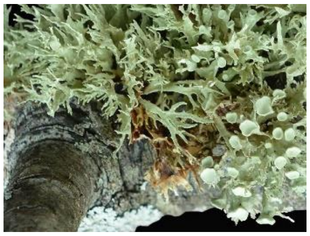
129

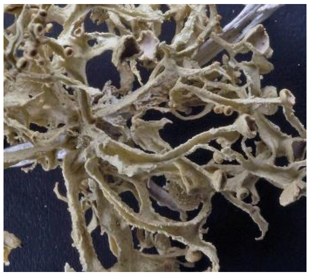
130

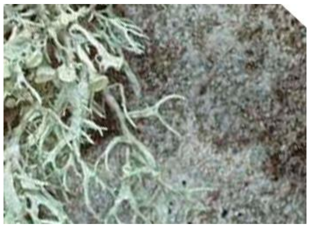
131

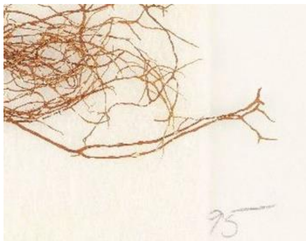

132

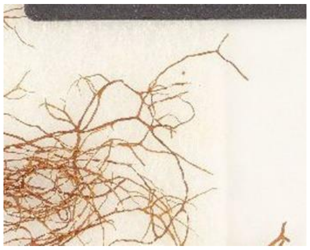
133

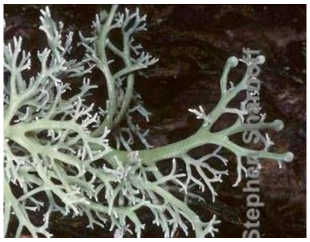
134

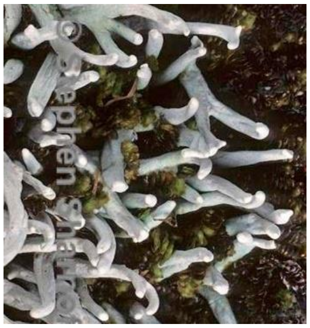
135

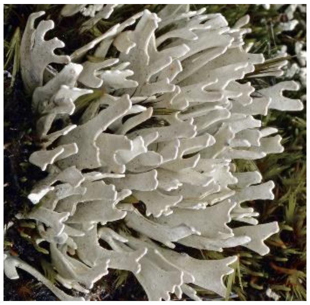
136

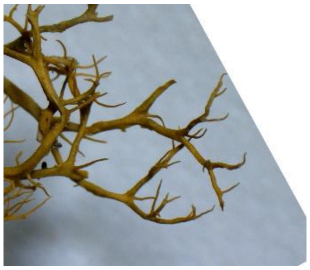
137

138 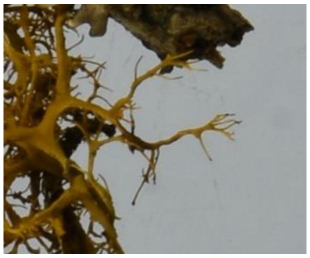 139 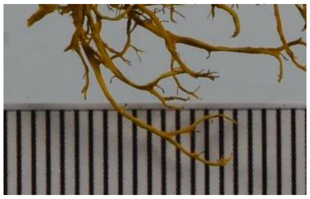

**Fig. S3.** The 140 images of five groups, i.e. microfoliose group, the *Daohugouthallus ciliiferus* fossil group, a long branches group, a wide-lobed group, and a fruticose group, used for geometric morphometric analysis and the order number in front of the image corresponds to the ID in Table S3.

**Table S1.** The photos information of 61 representative extant species and two fossil species in different orders of Lecanoromycetes used in the geometric morphometrics analysis

| **Sample information** | **Species** | **Family** | **Order** |
|---|---|---|---|
| CNALH Image Library (2 photo) | *Cladonia amaurocraea* | Cladoniaceae | Lecanorales |
| CNALH Image Library (1 photo) | *Cl. arbuscula* | Cladoniaceae | Lecanorales |
| HMAS-L 7879, Hubei, China (2 photos) | *Cl. furcata* | Cladoniaceae | Lecanorales |
| HMAS-L 122801, Jilin, China (2 photos) | *Cl. furcata* | Cladoniaceae | Lecanorales |
| CNALH Image Library (2 photos) | *Cl. furcata* | Cladoniaceae | Lecanorales |
| HMAS-L 120346, Shaanxi, China | *Cl. rangiferina* | Cladoniaceae | Lecanorales |
| HMAS-L 9403, Heilongjiang, China | *Evernia esorediosa* | Parmeliaceae | Lecanorales |
| HMAS-L 9405, Inner Mongolia, China | *E. esorediosa* | Parmeliaceae | Lecanorales |
| CNALH Image Library (2 photos) | *Hypotrachyna alectorialiorum* | Parmeliaceae | Lecanorales |
| CNALH Image Library (1 photos) | *H. britannica* | Parmeliaceae | Lecanorales |
| CNALH Image Library (3 photos) | *H. catawbiense* | Parmeliaceae | Lecanorales |
| HMAS-L 8322, Yunnan, China | *H. cirrhatum* | Parmeliaceae | Lecanorales |
| HMAS-L 8295, Guizhou, China | *H. cirrhatum* | Parmeliaceae | Lecanorales |
| HMAS-L 8319, Tibet, China | *H. cirrhatum* | Parmeliaceae | Lecanorales |
| HMAS-L 8339, Yunnan, China | *H. cirrhatum* | Parmeliaceae | Lecanorales |
| HMAS-L 22474, Yunnan, China | *H. cirrhatum* | Parmeliaceae | Lecanorales |
| HMAS-L 113576, Yunnan, China(2 photos) | *H. cirrhatum* | Parmeliaceae | Lecanorales |
| CNALH Image Library (2 photos) | *H. croceopustulata* | Parmeliaceae | Lecanorales |
| CNALH Image Library (3 photos) | *H. goiasii* | Parmeliaceae | Lecanorales |
| HMAS-L 8085, Yunnan, China | *H. nepalense* | Parmeliaceae | Lecanorales |
| HMAS-L 8269, Tibet, China | *H. rhizodendroideum* | Parmeliaceae | Lecanorales |
| HMAS-L 80658, Yunnan, China | *H. rhizodendroideum* | Parmeliaceae | Lecanorales |
| CNALH Image Library (2 photos) | *H. sinuosa* | Parmeliaceae | Lecanorales |
| CNALH Image Library (1 photos) | *H. vexans* | Parmeliaceae | Lecanorales |
| 24855, *Hypogymnia* Media Gallery | *Hypogymnia arcuata* | Parmeliaceae | Lecanorales |
| 31157, *Hypogymnia* Media Gallery | *H. fragillima* | Parmeliaceae | Lecanorales |
| dor-l24_1, *Hypogymnia* Media Gallery | *H. fragillima* | Parmeliaceae | Lecanorales |
| 31296, *Hypogymnia* Media Gallery | *H. physodes* | Parmeliaceae | Lecanorales |
| HMAS-L 81529, Xinjiang, China | *H. physodes* | Parmeliaceae | Lecanorales |
| HMAS-L 81350, Inner Mongolia, China | *H. vittata* | Parmeliaceae | Lecanorales |
| 25623, *Hypogymnia* Media Gallery | *H. vittata* | Parmeliaceae | Lecanorales |
| GZG_BST_21949 (fossil) | - | Parmeliaceae | Lecanorales |

| | | | |
|---|---|---|---|
| BitterfeldJörg Wunderlich Amber Collection F781 (fossil) | - | Parmeliaceae | Lecanorales |
| CNALH Image Library (2 photos) | *Lobaria anomala* | Lobariaceae | Peltigerales |
| CNALH Image Library (1 photos) | *L. hallii* | Lobariaceae | Peltigerales |
| CNALH Image Library (2 photos) | *L. kurokawae* | Lobariaceae | Peltigerales |
| CNALH Image Library (1 photos) | *L. linita* | Lobariaceae | Peltigerales |
| CNALH Image Library (1 photos) | *L. oregana* | Lobariaceae | Peltigerales |
| CNALH Image Library (1 photos) | *L. tenuis* | Lobariaceae | Peltigerales |
| CNALH Image Library (1 photos) | *Peltigera aphthosa* | Peltigeraceae | Peltigerales |
| CNALH Image Library (1 photos) | *P. castanea* | Peltigeraceae | Peltigerales |
| CNALH Image Library (3 photos) | *P. collina* | Peltigeraceae | Peltigerales |
| HMAS-L 13030, Inner Mongolia, China | *P. praetextata* | Peltigeraceae | Peltigerales |
| CNALH Image Library (1 photo) | *P. praetextata* | Peltigeraceae | Peltigerales |
| CNALH Image Library (1 photo) | *P. rufescens* | Peltigeraceae | Peltigerales |
| Colombia, Lucking sn_2766 (1 photo) | *Pannaria rubiginosa* | Pannariaceae | Peltigerales |
| Colombia, Lucking sn_2355 (1 photo) | *Pannaria rubiginosa* | Pannariaceae | Peltigerales |
| Colombia, Lucking sn_2719 (1 photo) | *Pannaria rubiginosa* | Pannariaceae | Peltigerales |
| Colombia, Lucking 41024a_4802 (1 photo) | *Pannaria rubiginosa* | Pannariaceae | Peltigerales |
| Colombia, Marin1648_6790 (1 photo) | *Coccocarpia stellata* | Coccocarpiaceae | Peltigerales |
| Costa Rica, Lucking (1 photo) | *Coccocarpia stellata* | Coccocarpiaceae | Peltigerales |
| HMAS-L 129385, Inner Mongolia, China | *Physcia adscendens* | Physciaceae | Caliciales |
| CNALH Image Library (2 photos) | *P. adscendens* | Physciaceae | Caliciales |
| CNALH Image Library (1 photos) | *P. aipolia* | Physciaceae | Caliciales |
| CNALH Image Library (2 photos) | *P. americana* | Physciaceae | Caliciales |
| CNALH Image Library (1 photos) | *P. biziana* | Physciaceae | Caliciales |
| CNALH Image Library (3 photos) | *P. caesia* | Physciaceae | Caliciales |
| CNALH Image Library (1 photos) | *P. halei* | Physciaceae | Caliciales |
| CNALH Image Library (2 photos) | *P. magnussonii* | Physciaceae | Caliciales |
| CNALH Image Library (4 photos) | *P. phaea* | Physciaceae | Caliciales |
| CNALH Image Library (1 photos) | *P. rolfii* | Physciaceae | Caliciales |
| CNALH Image Library (2 photos) | *P. stellaris* | Physciaceae | Caliciales |
| CNALH Image Library (1 photos) | *P. subalbinea* | Physciaceae | Caliciales |
| HMAS-L 110982, Xinjiang, China | *P. tenella* | Physciaceae | Caliciales |
| HMAS-L 110984, Xinjiang, China | *P. tenella* | Physciaceae | Caliciales |
| CNALH Image Library (1 photos) | *P. thomsoniana* | Physciaceae | Caliciales |
| CNALH Image Library (1 photos) | *Ramalina bajacalifornica* | Ramalinaceae | Lecanorales |
| CNALH Image Library (1 photos) | *R. calicaris* var. *fastigiata* | Ramalinaceae | Lecanorales |
| CNALH Image Library (1 photos) | *R. camptospora* | Ramalinaceae | Lecanorales |

| | | | |
|---|---|---|---|
| CNALH Image Library (1 photos) | *R. culbersoniorum* | Ramalinaceae | Lecanorales |
| CNALH Image Library (1 photos) | *R. darwiniana* var. *darwiniana* | Ramalinaceae | Lecanorales |
| CNALH Image Library (1 photos) | *R. dilacerata* | Ramalinaceae | Lecanorales |
| CNALH Image Library (2 photos) | *R. pacifica* | Ramalinaceae | Lecanorales |
| CNALH Image Library (1 photo) | *Sphaerophorus globosus* | Sphaerophoraceae | Lecanorales |
| CNALH Image Library (2 photos) | *Sticta beauvoisii* | Lobariaceae | Peltigerales |
| CNALH Image Library (1 photos) | *S. limbata* | Lobariaceae | Peltigerales |
| CNALH Image Library (1 photos) | *S. plumbicolor* | Lobariaceae | Peltigerales |
| CNALH Image Library (1 photos) | *S. xanthotropa* | Lobariaceae | Peltigerales |
| CNALH Image Library (1 photos) | *Siphula ceratites* | Icmadophilaceae | Pertusariales |
| Colombia, Lucking sn_9412 (1 photo) | *S. pteruloides* | Icmadophilaceae | Pertusariales |
| HMAS-L 504, Heilongjiang, China | *Teloschistes flavicans* | Teloschistaceae | Teloschistales |
| HMAS-L 104547, Hubei, China | *T. flavicans* | Teloschistaceae | Teloschistales |
| HMAS-L 104536, Sichuan, China | *T. flavicans* | Teloschistaceae | Teloschistales |
| Colombia, Lucking 41014_4736 (1 photo) | *Umbilicaria polyphylla* | Umbilicariaceae | Umbilicariales |
| Colombia, Lucking 41014_4737 (1 photo) | *Umbilicaria polyphylla* | Umbilicariaceae | Umbilicariales |
| Colombia, Lucking 41014_4738 (1 photo) | *Umbilicaria polyphylla* | Umbilicariaceae | Umbilicariales |
| Colombia, Lucking 41014_4739 (1 photo) | *Umbilicaria polyphylla* | Umbilicariaceae | Umbilicariales |
| Colombia, Lucking 41014_4743 (1 photo) | *Umbilicaria polyphylla* | Umbilicariaceae | Umbilicariales |

**Table S2.** The geometric morphometric analysis resulted in the variance and cumulative values for all the principal components

| **Principal component** | **Eigenvalues** | **Variance%** | **Cumulative %** |
|---|---|---|---|
| **1** | **0.03200622** | **24.376** | **24.376** |
| **2** | **0.02319641** | **17.666** | **42.042** |
| **3** | **0.01454791** | **11.079** | **53.121** |
| **4** | **0.01046854** | **7.973** | **61.094** |
| 5 | 0.0073343 | 5.586 | 66.679 |
| 6 | 0.006 | 4.57 | 71.249 |
| 7 | 0.00472941 | 3.602 | 74.851 |
| 8 | 0.00444899 | 3.388 | 78.239 |
| 9 | 0.00338804 | 2.58 | 80.819 |
| 10 | 0.00291705 | 2.222 | 83.041 |
| 11 | 0.00246823 | 1.88 | 84.921 |
| 12 | 0.00222106 | 1.692 | 86.612 |
| 13 | 0.0021387 | 1.629 | 88.241 |
| 14 | 0.00177234 | 1.35 | 89.591 |
| 15 | 0.00167563 | 1.276 | 90.867 |
| 16 | 0.00148536 | 1.131 | 91.998 |
| 17 | 0.00107806 | 0.821 | 92.819 |
| 18 | 0.00100053 | 0.762 | 93.581 |
| 19 | 0.00087003 | 0.663 | 94.244 |
| 20 | 0.00070831 | 0.539 | 94.783 |
| 21 | 0.00065028 | 0.495 | 95.279 |
| 22 | 0.00060754 | 0.463 | 95.741 |
| 23 | 0.00051279 | 0.391 | 96.132 |
| 24 | 0.00046675 | 0.355 | 96.487 |
| 25 | 0.00043885 | 0.334 | 96.821 |
| 26 | 0.00042747 | 0.326 | 97.147 |
| 27 | 0.00034762 | 0.265 | 97.412 |
| 28 | 0.00029405 | 0.224 | 97.636 |
| 29 | 0.00026663 | 0.203 | 97.839 |
| 30 | 0.00024143 | 0.184 | 98.023 |
| 31 | 0.00020254 | 0.154 | 98.177 |
| 32 | 0.00019285 | 0.147 | 98.324 |
| 33 | 0.00018533 | 0.141 | 98.465 |
| 34 | 0.00016945 | 0.129 | 98.594 |
| 35 | 0.0001465 | 0.112 | 98.706 |
| 36 | 0.00013846 | 0.105 | 98.811 |
| 37 | 0.00012771 | 0.097 | 98.908 |
| 38 | 0.00011912 | 0.091 | 98.999 |
| 39 | 0.0001087 | 0.083 | 99.082 |
| 40 | 0.00010072 | 0.077 | 99.158 |
| 41 | 0.00008652 | 0.066 | 99.224 |
| 42 | 0.00008538 | 0.065 | 99.289 |

| | | | |
|---|---|---|---|
| 43 | 0.00007168 | 0.055 | 99.344 |
| 44 | 0.00006718 | 0.051 | 99.395 |
| 45 | 0.00005783 | 0.044 | 99.439 |
| 46 | 0.00005578 | 0.042 | 99.482 |
| 47 | 0.00004892 | 0.037 | 99.519 |
| 48 | 0.00004683 | 0.036 | 99.555 |
| 49 | 0.00004168 | 0.032 | 99.586 |
| 50 | 0.00003921 | 0.03 | 99.616 |
| 51 | 0.00003592 | 0.027 | 99.644 |
| 52 | 0.00003432 | 0.026 | 99.67 |
| 53 | 0.00003169 | 0.024 | 99.694 |
| 54 | 0.00002899 | 0.022 | 99.716 |
| 55 | 0.00002772 | 0.021 | 99.737 |
| 56 | 0.00002557 | 0.019 | 99.756 |
| 57 | 0.00002491 | 0.019 | 99.775 |
| 58 | 0.0000216 | 0.016 | 99.792 |
| 59 | 0.00002129 | 0.016 | 99.808 |
| 60 | 0.00001817 | 0.014 | 99.822 |
| 61 | 0.00001739 | 0.013 | 99.835 |
| 62 | 0.00001461 | 0.011 | 99.846 |
| 63 | 0.00001424 | 0.011 | 99.857 |
| 64 | 0.00001318 | 0.01 | 99.867 |
| 65 | 0.00001241 | 0.009 | 99.877 |
| 66 | 0.00001202 | 0.009 | 99.886 |
| 67 | 0.00001102 | 0.008 | 99.894 |
| 68 | 0.00001024 | 0.008 | 99.902 |
| 69 | 0.00000998 | 0.008 | 99.91 |
| 70 | 0.00000892 | 0.007 | 99.916 |
| 71 | 0.00000794 | 0.006 | 99.922 |
| 72 | 0.00000741 | 0.006 | 99.928 |
| 73 | 0.00000727 | 0.006 | 99.934 |
| 74 | 0.00000635 | 0.005 | 99.938 |
| 75 | 0.00000598 | 0.005 | 99.943 |
| 76 | 0.00000591 | 0.005 | 99.948 |
| 77 | 0.00000553 | 0.004 | 99.952 |
| 78 | 0.00000491 | 0.004 | 99.955 |
| 79 | 0.00000457 | 0.003 | 99.959 |
| 80 | 0.00000429 | 0.003 | 99.962 |
| 81 | 0.0000041 | 0.003 | 99.965 |
| 82 | 0.0000037 | 0.003 | 99.968 |
| 83 | 0.00000348 | 0.003 | 99.971 |
| 84 | 0.00000318 | 0.002 | 99.973 |
| 85 | 0.00000307 | 0.002 | 99.976 |
| 86 | 0.00000271 | 0.002 | 99.978 |
| 87 | 0.00000253 | 0.002 | 99.98 |
| 88 | 0.00000252 | 0.002 | 99.981 |

| | | | |
|---|---|---|---|
| 89 | 0.00000245 | 0.002 | 99.983 |
| 90 | 0.00000227 | 0.002 | 99.985 |
| 91 | 0.00000208 | 0.002 | 99.987 |
| 92 | 0.00000197 | 0.002 | 99.988 |
| 93 | 0.00000167 | 0.001 | 99.989 |
| 94 | 0.00000154 | 0.001 | 99.991 |
| 95 | 0.00000148 | 0.001 | 99.992 |
| 96 | 0.00000131 | 0.001 | 99.993 |
| 97 | 0.0000013 | 0.001 | 99.994 |
| 98 | 0.00000112 | 0.001 | 99.995 |
| 99 | 0.000001 | 0.001 | 99.995 |
| 100 | 0.00000085 | 0.001 | 99.996 |
| 101 | 0.00000074 | 0.001 | 99.997 |
| 102 | 0.0000006 | 0 | 99.997 |
| 103 | 0.00000057 | 0 | 99.997 |
| 104 | 0.00000054 | 0 | 99.998 |
| 105 | 0.00000053 | 0 | 99.998 |
| 106 | 0.00000041 | 0 | 99.999 |
| 107 | 0.00000038 | 0 | 99.999 |
| 108 | 0.00000032 | 0 | 99.999 |
| 109 | 0.0000003 | 0 | 99.999 |
| 110 | 0.00000021 | 0 | 99.999 |
| 111 | 0.00000019 | 0 | 100 |
| 112 | 0.00000019 | 0 | 100 |
| 113 | 0.00000014 | 0 | 100 |
| 114 | 0.00000012 | 0 | 100 |
| 115 | 0.00000005 | 0 | 100 |
| 116 | 0.00000003 | 0 | 100 |

**Table S3.** The data set of five groups divided according to thallus growth types used in the geometric morphometrics analysis

| ID | Photo name | Group |
|---|---|---|
| **0** | FO-C-Phy-*Physcia adscendens* HMAS129385 | 1 |
| **1** | FO-C-Phy-*Physcia adscendens* | 1 |
| **2** | FO-C-Phy-*Physcia adscenden*s-1 | 1 |
| **3** | FO-C-Phy-*Physcia aipolia*_9 | 1 |
| **4** | FO-C-Phy-*Physcia americana*_1 | 1 |
| **5** | FO-C-Phy-*Physcia americana*_2 | 1 |
| **6** | FO-C-Phy-*Physcia biziana*_9 | 1 |
| **7** | FO-C-Phy-*Physcia caesia*-1 | 1 |
| **8** | FO-C-Phy-*Physcia caesia*-2 - 1 | 1 |
| **9** | FO-C-Phy-*Physcia caesia*-2 | 1 |
| **10** | FO-C-Phy-*Physcia halei*_3 | 1 |
| **11** | FO-C-Phy-*Physcia magnussonii* | 1 |
| **12** | FO-C-Phy-*Physcia magnussonii*-1 | 1 |
| **13** | FO-C-Phy-*Physcia phaea_poss* | 1 |
| **14** | FO-C-Phy-*Physcia phaea*_poss-1 | 1 |
| **15** | FO-C-Phy-*Physcia phaea*_poss-2 | 1 |
| **16** | FO-C-Phy-*Physcia phaea*_poss-3 | 1 |
| **17** | FO-C-Phy-*Physcia rolfii* | 1 |
| **18** | FO-C-Phy-*Physcia stellaris* | 1 |
| **19** | FO-C-Phy-*Physcia stellaris*-1 | 1 |
| **20** | FO-C-Phy-*Physcia subalbinea*-1 | 1 |
| **21** | FO-C-Phy-*Physcia tenella* HMAS110982 | 1 |
| **22** | FO-C-Phy-*Physcia tenella* HMAS110984 | 1 |
| **23** | FO-C-Phy-*Physcia thomsoniana* | 1 |
| **24** | FO-P-Coc-*Coccocarpia stellata* 1648_6790 | 1 |
| **25** | FO-P-Coc-*Coccocarpia stellata* | 1 |
| **26** | FO-P-Pan-*Pannaria rubiginosa* 2766 | 1 |
| **27** | FO-P-Pan-*Pannaria rubiginosa* 2355 | 1 |
| **28** | FO-P-Pan-*Pannaria rubiginosa* 2719 | 1 |
| **29** | FO-P-Pan-*Pannaria rubiginosa* 41024a_4802 | 1 |
| **30** | FO-L-Dao-*Daohugouthallus ciliiferus*-1 | 2 |
| **31** | FO-L-Dao-*Daohugouthallus ciliiferus*-2 | 2 |
| **32** | FO-L-Dao-*Daohugouthallus ciliiferus*-3 | 2 |
| **33** | FO-L-Dao-*Daohugouthallus ciliiferus*-4 | 2 |
| **34** | FO-L-Dao-*Daohugouthallus ciliiferus*-5 | 2 |
| **35** | FO-L-Dao-*Daohugouthallus ciliiferus*-B0476P -1 | 2 |
| **36** | FO-L-Dao-*Daohugouthallus ciliiferus*-B0476P | 2 |

| | | |
|---|---|---|
| **37** | FO-L-Dao-*Daohugouthallus ciliiferus*-NN2019001-1 | 2 |
| **38** | FO-L-Dao-*Daohugouthallus ciliiferus*-NN2019001-2 | 2 |
| **39** | FO-L-Dao-*Daohugouthallus ciliiferus*-NN2019001-3 | 2 |
| **40** | FO-L-Dao-*Daohugouthallus ciliiferus*-NN2019001-4 | 2 |
| **41** | FO-L-Dao-*Daohugouthallus ciliiferus*-NN2019001-5 | 2 |
| **42** | FO-L-Dao-*Daohugouthallus ciliiferus*-NN2019001-6 | 2 |
| **43** | FO-L-Dao-*Daohugouthallus ciliiferus*-NN2019001-7 | 2 |
| **44** | FO-L-Dao-*Daohugouthallus ciliiferus*-NN2019001-8 | 2 |
| **45** | FO-L-Dao-*Daohugouthallus ciliiferus*-new1-1 | 2 |
| **46** | FO-L-Dao-*Daohugouthallus ciliiferus*-new1-2 | 2 |
| **47** | FO-L-Dao-*Daohugouthallus ciliiferus*-new1-3 | 2 |
| **48** | FO-L-Dao-*Daohugouthallus ciliiferus*-new1-4 | 2 |
| **49** | FO-L-Dao-*Daohugouthallus ciliiferus*-new1-5 | 2 |
| **50** | FO-L-Dao-*Daohugouthallus ciliiferus*-new1-6 | 2 |
| **51** | FO-L-Dao-*Daohugouthallus ciliiferus*-new1-7 | 2 |
| **52** | FO-L-Dao-*Daohugouthallus ciliiferus*-new1-8 | 2 |
| **53** | FO-L-Dao-*Daohugouthallus ciliiferus*-new1-9 | 2 |
| **54** | FO-L-Dao-*Daohugouthallus ciliiferus*-new1-10 | 2 |
| **55** | FO-L-P-GZG BST 21949 | 3 |
| **56** | FO-L-Par-*Hypogymnia arcuata*-24855 | 3 |
| **57** | FO-L-Par-*Hypogymnia fragillima*-31157 | 3 |
| **58** | FO-L-Par-*Hypogymnia fragillima*-dorl24_1 | 3 |
| **59** | FO-L-Par-*Hypogymnia physodes* HMAS81529 | 3 |
| **60** | FO-L-Par-*Hypogymnia physodes*-31296 | 3 |
| **61** | FO-L-Par-*Hypogymnia vittata* HMAS81350 | 3 |
| **62** | FO-L-Par-*Hypogymnia vittata*-25623 | 3 |
| **63** | FO-L-Par-*Hypotrachyna alectorialiorum*-1 | 3 |
| **64** | FO-L-Par-*Hypotrachyna alectorialiorum*-2 | 3 |
| **65** | FO-L-Par-*Hypotrachyna britannica* | 3 |
| **66** | FO-L-Par-*Hypotrachyna catawbiense*_1-1 | 3 |
| **67** | FO-L-Par-*Hypotrachyna catawbiense*_1-2 | 3 |
| **68** | FO-L-Par-*Hypotrachyna catawbiense*_1-3 | 3 |
| **69** | FO-L-Par-*Hypotrachyna cirrhatum* HMAS8295 | 3 |
| **70** | FO-L-Par-*Hypotrachyna cirrhatum* HMAS8319 | 3 |
| **71** | FO-L-Par-*Hypotrachyna cirrhatum* HMAS8322 | 3 |
| **72** | FO-L-Par-*Hypotrachyna cirrhatum* HMAS8339 | 3 |
| **73** | FO-L-Par-*Hypotrachyna cirrhatum* HMAS22474 | 3 |
| **74** | FO-L-Par-*Hypotrachyna cirrhatum* HMAS113576 | 3 |
| **75** | FO-L-Par-*Hypotrachyna cirrhatum* HMAS113576-1 | 3 |
| **76** | FO-L-Par-*Hypotrachyna croceopustulata* | 3 |
| **77** | FO-L-Par-*Hypotrachyna croceopustulata*-1 | 3 |

| | | |
|---|---|---|
| **78** | FO-L-Par-*Hypotrachyna goiasii*-1 | 3 |
| **79** | FO-L-Par-*Hypotrachyna goiasii*-2 | 3 |
| **80** | FO-L-Par-*Hypotrachyna goiasii*-3 | 3 |
| **81** | FO-L-Par-*Hypotrachyna nepalense* HMAS8085 | 3 |
| **82** | FO-L-Par-*Hypotrachyna rhizodendroideum* HMAS8269 | 3 |
| **83** | FO-L-Par-*Hypotrachyna rhizodendroideum* HMAS80658 | 3 |
| **84** | FO-L-Par-*Hypotrachyna sinuosa* | 3 |
| **85** | FO-L-Par-*Hypotrachyna sinuosa*-1 | 3 |
| **86** | FO-L-Par-*Hypotrachyna vexans*-1 | 3 |
| **87** | FO-P-Lob-*Lobaria anomala* | 4 |
| **88** | FO-P-Lob-*Lobaria anomala*_7 | 4 |
| **89** | FO-P-Lob-*Lobaria hallii*_8 | 4 |
| **90** | FO-P-Lob-*Lobaria kurokawae*-van-herk_web | 4 |
| **91** | FO-P-Lob-*Lobaria kurokawae*-van-herk-1 | 4 |
| **92** | FO-P-Lob-*Lobaria linita*_4 | 4 |
| **93** | FO-P-Lob-*Lobaria oregana*_9 | 4 |
| **94** | FO-P-Lob-*Lobaria tenuis* | 4 |
| **95** | FO-P-Lob-*Sticta beauvoisii* | 4 |
| **96** | FO-P-Lob-*Sticta beauvoisii*_3_ | 4 |
| **97** | FO-P-Lob-*Sticta limbata* | 4 |
| **98** | FO-P-Lob-*Sticta plumbicolor*-hawaii-3108279 | 4 |
| **99** | FO-P-Lob-*Sticta xanthotropa*-french-guiana-5914774 | 4 |
| **100** | FO-P-Pel-*Peltigera aphthosa*_3 | 4 |
| **101** | FO-P-Pel-*Peltigera castanea* | 4 |
| **102** | FO-P-Pel-*Peltigera collina* | 4 |
| **103** | FO-P-Pel-*Peltigera collina*-1 | 4 |
| **104** | FO-P-Pel-*Peltigera collina*-2 | 4 |
| **105** | FO-P-Pel-*Peltigera praetextata* HMAS13030 | 4 |
| **106** | FO-P-Pel-*Peltigera praetextata* | 4 |
| **107** | FO-P-Pel-*Peltigera rufescens*_cfr | 4 |
| **108** | FO-U-Umb-*Umbilicaria polyphylla* 41014_4736 | 4 |
| **109** | FO-U-Umb-*Umbilicaria polyphylla* 41014_4737 | 4 |
| **110** | FO-U-Umb-*Umbilicaria polyphylla* 41014_4738 | 4 |
| **111** | FO-U-Umb-*Umbilicaria polyphylla* 41014_4739 | 4 |
| **112** | FO-U-Umb-*Umbilicaria polyphylla* 41014_4743 | 4 |
| **113** | FR-L-Cla-*Cladonia amaurocraea*-1 | 5 |
| **114** | FR-L-Cla-*Cladonia amaurocraea*-1-1 | 5 |
| **115** | FR-L-Cla-*Cladonia arbuscula* | 5 |
| **116** | FR-L-Cla-*Cladonia furcata* HMAS7879 | 5 |
| **117** | FR-L-Cla-*Cladonia furcata* HMAS7879-1 | 5 |
| **118** | FR-L-Cla-*Cladonia furcata* HMAS122801 | 5 |
| **119** | FR-L-Cla-*Cladonia furcata* HMAS122801-1 | 5 |

| | | |
|---|---|---|
| **120** | FR-L-Cla-*Cladonia furcata*-1 | 5 |
| **121** | FR-L-Cla-*Cladonia furcata*-1-1 | 5 |
| **122** | FR-L-Cla-*Cladonia rangiferina* HMAS120346 | 5 |
| **123** | FR-L-P-BitterfeldJörg Wunderlich Amber Collection F781 | 5 |
| **124** | FR-L-Par-*Evernia esorediosa* HMAS9403 | 5 |
| **125** | FR-L-Par-*Evernia esorediosa* HMAS9405 | 5 |
| **126** | FR-L-Ram-*Ramalina bajacalifornica*_Sharn_1557251388 | 5 |
| **127** | FR-L-Ram-*Ramalina calicaris* var. *fastigiata* FH00511819 | 5 |
| **128** | FR-L-Ram-*Ramalina camptospora* Nyl. | 5 |
| **129** | FR-L-Ram-*Ramalina culbersoniorum*_1550624964 | 5 |
| **130** | FR-L-Ram-*Ramalina darwiniana* var. *darwiniana* | 5 |
| **131** | FR-L-Ram-*Ramalina dilacerata*_2 | 5 |
| **132** | FR-L-Ram-*Ramalina pacifica* Asahina | 5 |
| **133** | FR-L-Ram-*Ramalina pacifica* Asahina-1 | 5 |
| **134** | FR-L-Sph-*Sphaerophorus* globosus-1 | 5 |
| **135** | FR-P-Icm-*Siphula ceratites*_3 | 5 |
| **136** | FR-P-Icm-*Siphula pteruloides* 9412 | 5 |
| **137** | FR-T-Tel-*Teloschistes flavicans* HMAS504 | 5 |
| **138** | FR-T-Tel-*Teloschistes flavicans* HMAS547 | 5 |
| **139** | FR-T-Tel-*Teloschistes flavicans* HMAS104536 | 5 |

Notes: The photo name is listed as ‘thallus growth type-the first letter of Order name-the first three letters of Family name-species name–specimen No. FO=Foliose, FR=fruticose.
